# Paralogue-selective degradation of the lysine acetyltransferase EP300

**DOI:** 10.1101/2024.05.03.592353

**Authors:** Xuemin Chen, McKenna C. Crawford, Ying Xiong, Anver Basha Shaik, Kiall F. Suazo, Manini S. Penkalapati, Joycelyn H. Williams, Thorkell Andressen, Rolf E. Swenson, Jordan L. Meier

## Abstract

The transcriptional coactivators EP300 and CREBBP are critical regulators of gene expression that share high sequence identity but exhibit non-redundant functions in basal and pathological contexts. Here, we report the development of a bifunctional small molecule, MC-1, capable of selectively degrading EP300 over CREBBP. Using a potent aminopyridine-based inhibitor of the EP300/CREBBP catalytic domain in combination with a VHL ligand, we demonstrate that MC-1 preferentially degrades EP300 in a proteasome-dependent manner. Mechanistic studies reveal that selective degradation cannot be predicted solely by target engagement or ternary complex formation, suggesting additional factors govern paralogue-specific degradation. MC-1 inhibits cell proliferation in a subset of cancer cell lines and provides a new tool to investigate the non-catalytic functions of EP300 and CREBBP. Our findings expand the repertoire of EP300/CREBBP-targeting chemical probes and offer insights into the determinants of selective degradation of highly homologous proteins.

---

The paralogous transcriptional coactivators EP300 and CREBBP are critical regulators of transcription signaling in biology and disease.^1^ Within the nucleus, these high molecular weight (∼300 kDa) enzymes engage partner proteins through an array of protein-protein interaction motifs. Many of these proteins, including histones and transcription factors, serve as substrates for the catalytic histone acetyltransferase (HAT) domain embedded in the core of these proteins (Fig. 1). As central nodes in transcriptional signaling, EP300 and CREBBP have been implicated in both oncogenic and tumor suppressive function in cancer.^2^ However, despite their strong similarity, EP300 and CREBBP do not appear to be functionally redundant across all biological contexts. For instance, genetic disruption of EP300 alone is sufficient to impair the leukemogenicity of the AML1-ETO transcription factor^3^ and facilitate anti-tumor autoimmunity in T-regulatory cells.^4^ Conversely, CREBBP alone constitutes a unique non-oncogene dependency in EP300-deficient cancers^5^ and is required for hematopoietic stem cell self-renewal in mice.^6^ To date, paralogue-specific functions of EP300 and CREBBP have been almost exclusively studied using genetic methods.

**Figure 1.**
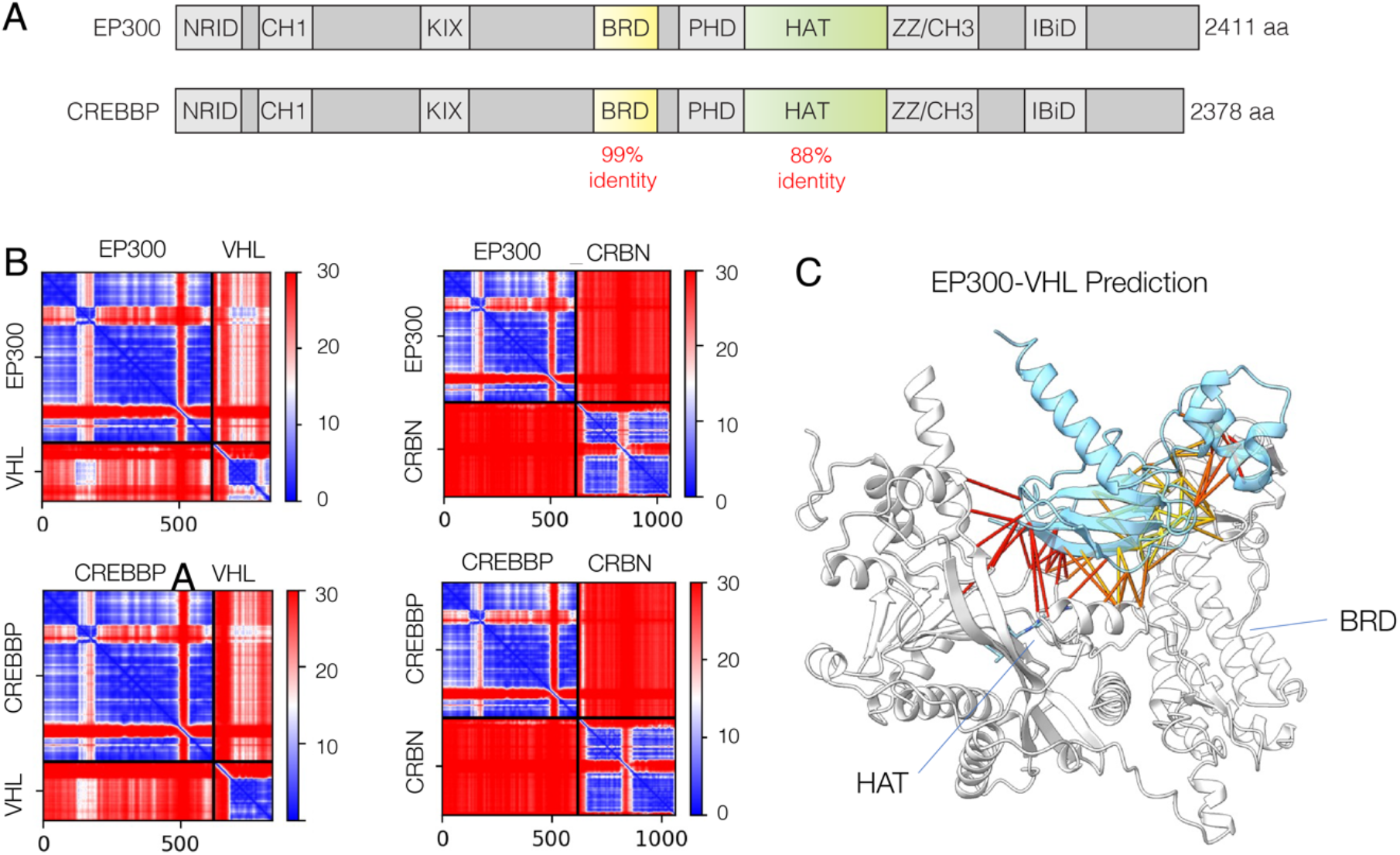
(A) Domain architecture of EP300 and CREBBP. The relative percent identity of two domains targeted by small molecules is specified. BRD = bromodomain, HAT = histone acetyltransferase domain. (B) AF-Multimer-generated 2D plot of predicted alignment error (PAE) from predictions of EP300-E3 ligase and CREBBP-E3 ligase complexes. CRBN = cereblon, VHL = von Hippel-Landau disease tumor suppressor. (C) AF-Multimer predicted structure of EP300 catalytic core (white) bound to VHL (blue). The majority of predicted interactions are with residues found in the EP300 HAT domain. All predictions were made using Colabfold.^19^

From a small molecule perspective, significant efforts have led to the development of potent and selective inhibitors of the EP300 and CREBBP HAT domain.^7, 8^ One of the most potent of these compounds is CPI-1612.^9, 10^ In biochemical assays CPI-1612 disrupts EP300 and CREBBP catalytic activity at half-maximal inhibitor concentrations of <0.5 nM and 2.9 nM, respectively (Table S1). Considering how this narrow selectivity window may be leveraged to study individual functions of EP300 and CREBBP, we were inspired by recent examples from the targeted protein degradation literature.^11^ This strategy entails the construction of bifunctional molecules that induce proximity between a target protein and an E3 ligase, which in turn triggers the target’s ubiquitinylation and proteasomal degradation. In one example, conversion of a dual CDK4/CDK6 ligand into a bifunctional enabled selective degradation of CDK6.^12^ Most EP300/CREBBP degraders reported to date engage the bromodomain and cause dual degradation of both paralogues.^13-16^ One degrader based on the HAT inhibitor A-485 has been shown to selectively degrade EP300 in neuroblastoma models.^17^ However, we found this molecule does not have significant activity in HAP-1 cells (Fig. S1), suggesting further exploration of paralogue-specific degradation strategies could be warranted.

To explore potential avenues for differentiating these closely related proteins, we began by examining the sequence conservation between the ligandable active sites of EP300 and CREBBP. EP300 and CREBBP’s HAT domains are 88% identical and 93% homologous, while their bromodomains share 97% identity and 99% homology (Fig. 1a). Superimposing the non-conserved residues in the HAT domain onto a crystal structure revealed the majority appear lie outside the small molecule binding site (Fig. S2). To assess the potential for bifunctional small molecules differentiate the two proteins based on these small changes, we used the Colabfold implementation of AlphaFold Multimer^18, 19^ to predict the structure of EP300 and CREBBP in complex with the most commonly recruited E3 ligases, VHL and CRBN. Different degrees of confidence were made in the predictions across the two proteins (Fig. 1b). Interestingly, the majority of predicted contacts were made with the HAT region, rather than the bromodomain (Fig. 1c). We take care not to overstate this observation, as recent analyses have indicated that current versions of AlphaFold are not useful for predicting the final conformation of E3 ligase/ternary complexes formed by induced proximity reagents.^20^ However, the potential for differential molecular recognition, together with slight biochemical selectivity of CPI-1612 for EP300, motivated us to explore this HAT ligand’s degrader capabilities.

A conjugatable analogue of CPI-1612 was synthesized via a convergent route (Scheme 1). The pyrazole amide was chosen as a suitable exit vector based on the published crystal structure of the CPI-1612/EP300 complex as well as a recent medicinal chemistry effort showing that alteration of this region is well-tolerated.^9, 21^ Briefly, ester **1** and amine **2** were coupled by nucleophilic displacement and purified to yield the enantiomerically pure (*R, S*) stereoisomer of **3**. Palladium coupling of boronate **4** with **5** provided aminopyridine **6**, which was further condensed with **3** using lithium hexamethyldisilazide to provide carboxylate **7**. Conventional amide coupling methods were then used to conjugate this precursor to a small panel of amine-containing analogues of known VHL and CRBN recruiters to afford bifunctional HAT domain ligands **8**-**19**.

**Scheme 1.**
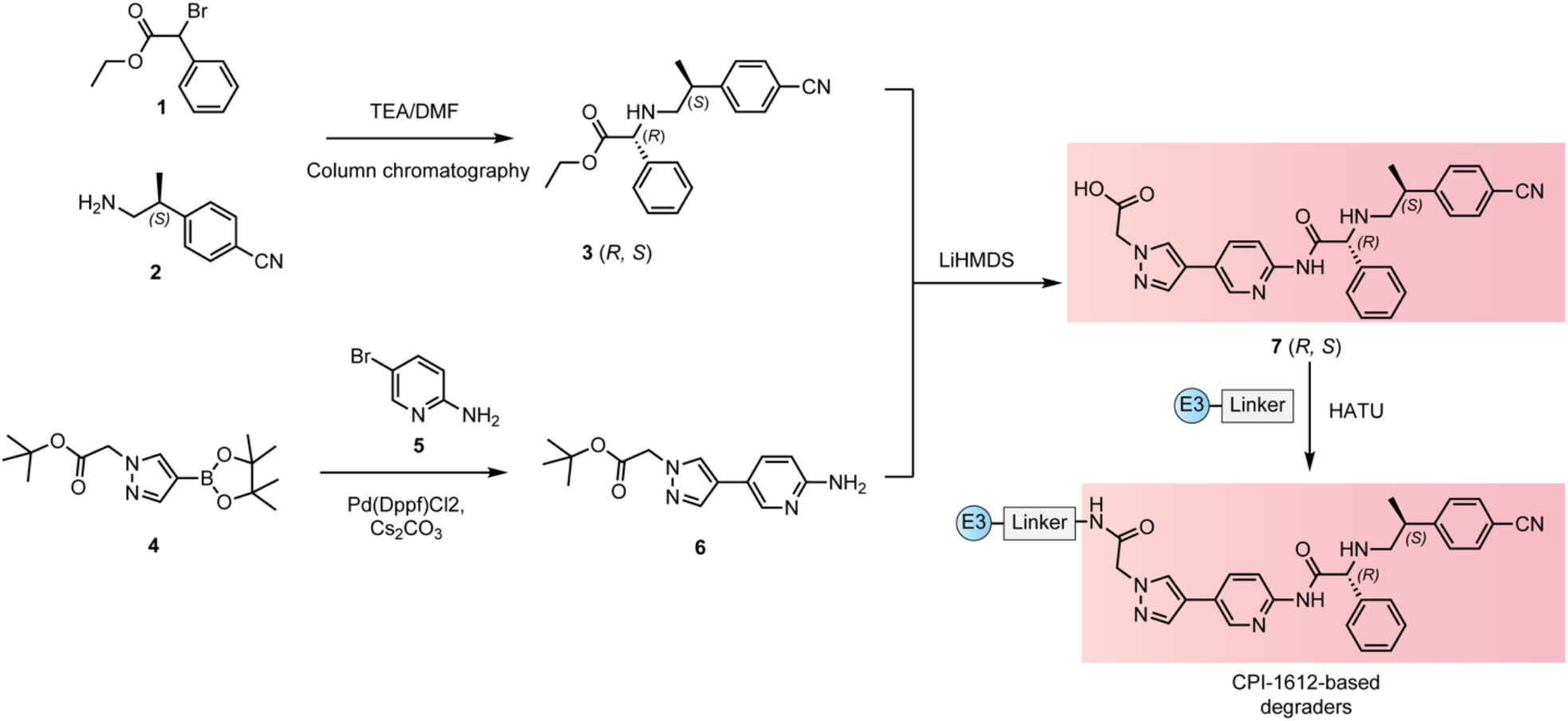
Synthesis of CPI-1612-based degrader molecules. Full synthetic details are provided in the Supporting Information.

As an initial test of our molecules to mediate EP300 and CREBBP degradation, we treated HAP-1 cells for approximately 24 hours with **8**-**19** across a concentration gradient of 500-10,000 nM (Fig. 2a). HAP-1 cells have previously been deployed in EP300/CREBBP degrader development.^13^ Furthermore, both VHL and CRBN-recruiting bifunctionals have been shown to be active in this model.^22^ Amongst VHL recruiting small molecules, C8-linked compound **10** was the most potent degrader, followed by shorter C5-linked **11** and pyridine **13** (Fig. 2b). All three molecules showed a preference for degradation of EP300 over CREBBP. This contrasted with control molecule dCBP-1 which degraded both paralogues (Fig. 2b).^13^ Amongst CRBN-based degraders, **15** was the most active and also appeared to show a slight preference for EP300. However, it triggered a visibly greater maximal degradation (D_max_) of CREBBP relative to compound **10** (Fig. S3). These results suggest that recruitment of E3 ligases to the EP300/CREBBP HAT domain can trigger preferential degradation of EP300.

**Figure 2.**
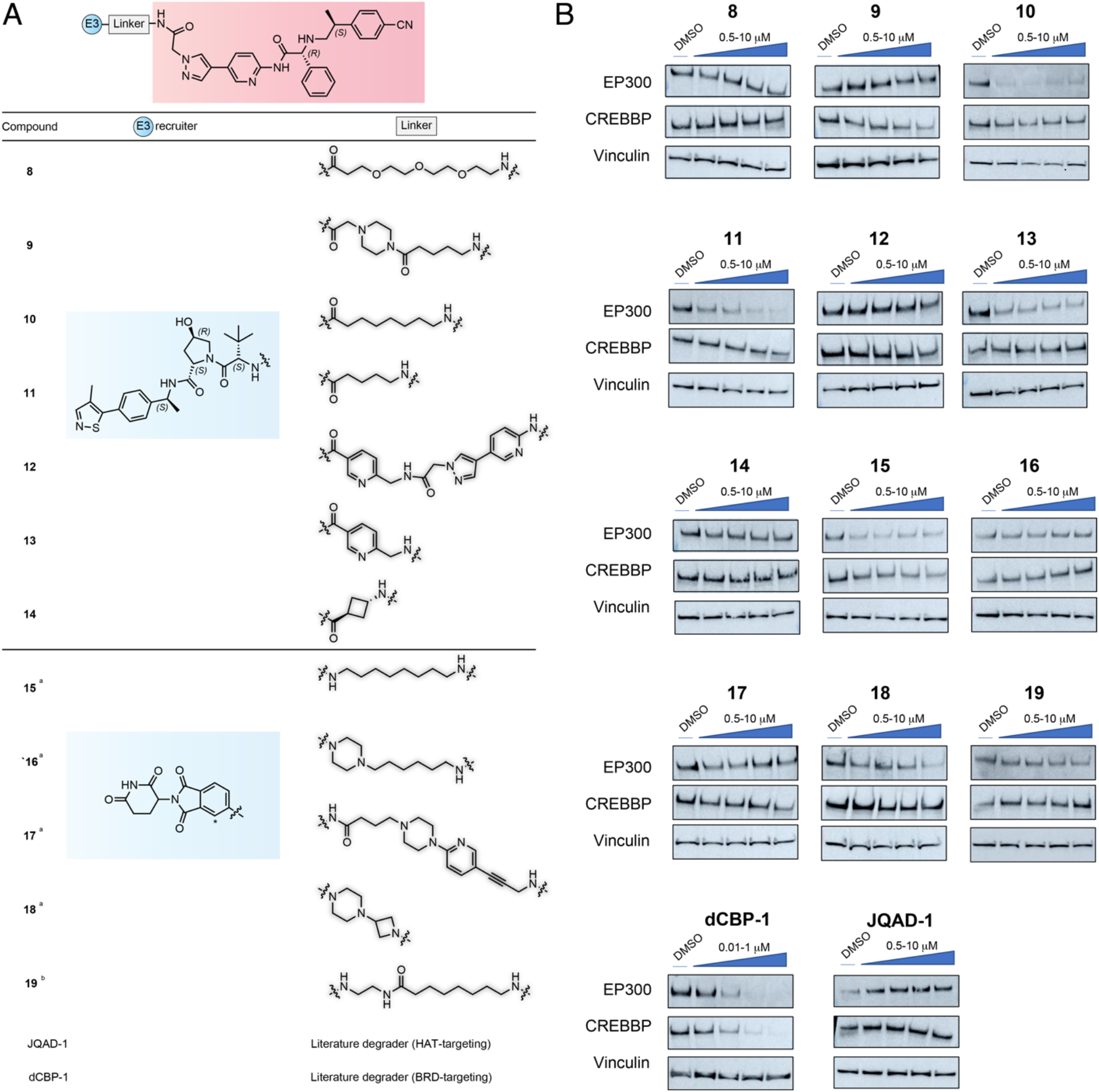
(A) CPI-1612-based degrader and molecules tested in this study. Compounds **8**-**14** use a VHL-recruiting ligand (blue), while compounds **15**-**19** feature a CRBN-recruiting ligand (blue). ^a^Molecule tested as a mixture of diastereomers at the phenylglycine stereocenter. ^b^Molecule linked CRBN ligand at the C2 position (denoted with *). Full structures are available in the Supporting Information. (B) Initial screen for modulation of EP300 and CREBBP levels at ∼24 h. Compounds **8**-**19** were tested at 0.5, 1, 5, and 10 μM. dCBP-1 was tested at 0.001, 0.01, 0.1 10 μM. JQAD-1 was tested at 0.1, 1, 5, and 10 μM. Results are representative of n = ≥2 biological replicates.

To assess the correlation between binding and EP300 degradation, we synthesized an affinity probe based on the CPI-1612 scaffold and performed competitive affinity pulldown experiments using lead degrader **10** (Fig. S4a). In this assay, ligands are pre-incubated with HeLa nuclear extracts and tested for their ability to block affinity capture of EP300 or CREBBP (Fig. S4b). The higher the concentration of free ligand required for competition, the weaker the binding. Compared to CPI-1612, we found that **10** only competed EP300 and CREBBP capture at higher concentrations (Fig. S4c-d, Fig. S5). Cell-based assays also indicate that CPI-1612 is a more potent inhibitor of EP300/CREBBP-catalyzed acetylation than **10** (Fig. S4e). This suggests degradation does not reflect selective preservation of CPI-1612’s intrinsic binding affinity for EP300 and CREBBP by compound **10**. Motivated by the results of our cellular experiment, we next tested whether using H3K18Ac as a readout for EP300/CREBBP occupancy could allow us to understand the relationship between target engagement and degradation more broadly across our panel. As inhibition of H3K18Ac by **8**-**19** provides a combined measure of cellular accumulation and EP300/CREBBP binding, we hypothesized it could be used to determine if HAT engagement is predictive of degradation. In contrast, observing efficient inhibition of H3K18Ac without degradation may reflect a molecule’s inability to form a productive protein-degrader-E3 ternary complex. Assayed at two concentrations, **10** and **11** were the most potent VHL-based inhibitors, while several molecules (**9, 12**-**14**) had little impact on acetylation (Fig. S6). CRBN-based compounds were overall more active as HAT inhibitors, in line with the cyclic imide warhead being more cell permeable.^23^ However, HAT engagement did not strictly correlate with degradation. This is most evident in the observation that **16** and **18** – the most potent inhibitors of H3K18Ac in our panel (Fig. S6) – did not affect levels of EP300 or CREBBP (Fig. 2b).

The finding that target engagement is necessary but not sufficient for EP300-selective degradation led us to further investigate ternary complex formation. For these studies we narrowed our focus to lead molecule **10**, which for convenience we refer to by the abbreviation “MC-1” (Fig. 3a). To monitor EP300-VHL interactions in real time in living cells, we employed a luciferase complementation assay based on split Nanoluc.^24^ Briefly, plasmid constructs were prepared in which the EP300 catalytic core consisting of the HAT and bromodomain (amino acids 1048-1665) was fused at the C-terminus to an inactive Nanoluc protein (LgBIT), and full length VHL was fused to a complementary fragment (SmBIT; Fig. 3b). Co-expression of these proteins does not reconstitute Nanoluc activity due to their low intrinsic affinity but can be stimulated by compounds capable of nucleating formation of a stable ternary complex. Following transfection of HEK-293T cells with C-terminal EP300-LgBIT and N-terminal SmBIT-VHL, the Nanoluc substrate furimazine was added and allowed to equilibrate. Addition of MC-1 induced a rapid increase in Nanoluc activity, consistent with ternary complex formation (Fig. 3c). Treatment of cells with CPI-1612 or vehicle (DMSO) did not cause a similar response. Performing an identical assay but replacing the EP300 with the CREBBP catalytic core also yielded data consistent with nucleation of a HAT-VHL interaction (Fig. S7). Further experimentation found EP300 degradation by MC-1 to be dose-dependent (Fig. 3d), time-dependent (Fig. 3e), and sensitive to ‘squelching’ by free VHL ligand (Fig. 3f) as well as CPI-1612 itself (Fig. 3g). These data provide further evidence that MC-1 functions via ternary complex formation, but indicate that this factor alone does not govern its preferential degradation of EP300 versus CREBBP.

**Figure 3.**
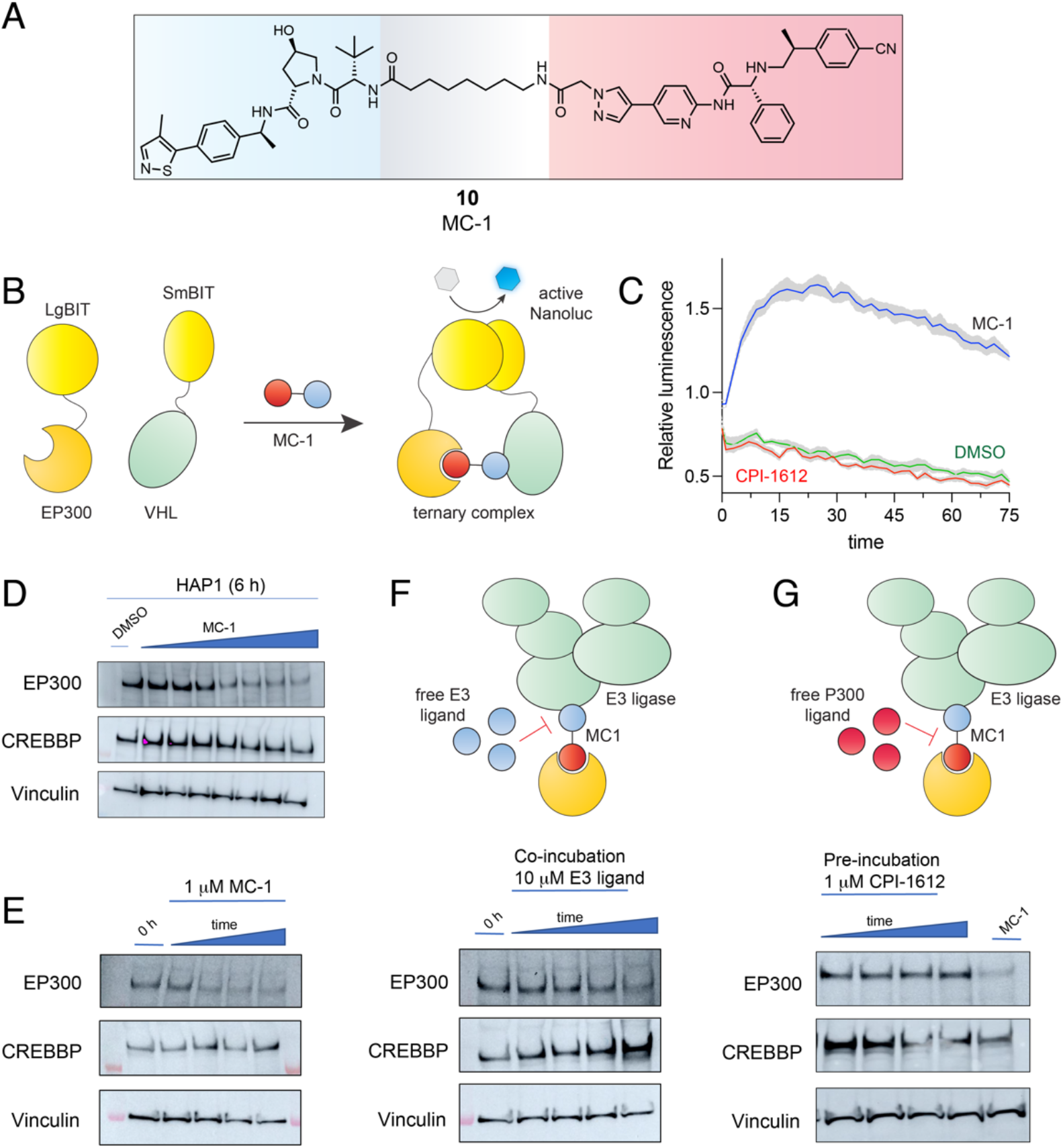
(A) Structure of lead degrader **10**, referred to in this manuscript as MC-1. (B) Schematic of NanoBiT assay for assessing small molecule-dependent EP300-VHL interaction (C) Measurement of EP300-VHL binding via NanoBiT assay. Averages of 4 replicates are plotted, shaded areas represent standard deviation of mean. (D) Dose-dependent degradation of EP300 at 6 h in HAP1 cells. Concentrations: 0.01, 0.05, 0.1, 0.25, 0.5, 1, 2.5 µM. (E) Time-dependent degradation of EP300 by 1 mM MC-1 in HAP1 cells. Time points: 0, 1, 3, 6, 24 h. (F) Testing the ability of free E3 ligase ligand (VH298, 10 mM) to block degradation of EP300 by MC-1 (1 mM) in HAP1 cells. Time points: 0, 1, 3, 6, 24 h. (G) Testing the ability of free HAT ligand (CPI-1612, 1 mM) to block degradation of EP300 by MC-1 (1 mM) in HAP1 cells. CPI-1612 was pre-incubated for 1 h before MC-1 addition. Time points (after MC-1 addition): 1, 3, 6, 24 h.

To investigate the broader impact of MC-1 on protein levels within cells, we conducted TMT-based proteomic profiling. These experiments compared HAP-1 cells treated with MC-1 (1 μM, 6 h) to control cells treated with vehicle DMSO. We also assessed dCBP-1 (250 nM, 6 h), using it as a validated probe known to degrade both EP300 and CREBBP. To maximize coverage, samples were extracted and subject to offline fractionation prior to tandem-mass tag labeling and pooled analysis. This strategy enabled the identification of >8,400 quantified proteins (Table S1). We defined a significant change in protein level as having a p-value less than 0.05 and a log2 fold change ratio of −0.5 (MC1-treated/DMSO-treated) and limited our reporter ion quantitation to unique peptides due to the high percent identity of EP300 and CREBBP. The number of PSMs assigned to CREBBP (19) and EP300 (9) are high enough to infer accurate quantitation in specifically comparing these two paralogues. Remarkably, this analysis revealed EP300 as the only HAT degraded upon MC-1 treatment (Fig. 4a). Applying similar criteria to dCBP-1 validated the ability of this probe to potently degrade both EP300 and CREBBP (Fig. 4b). Treatment of cells with MC-1 for 6 h also changed the abundance of additional proteins unrelated to histone acetylation, perhaps unsurprising given the centrality of EP300 to transcriptional homeostasis. The only one of these proteins exhibiting a log2 fold decrease >0.5 was the alternative splicing regulator NOVA2. Dual degrader dCBP-1 caused a larger proteomic perturbation than MC-1. Interestingly, dCBP-1 also regulated NOVA2. Given the disparate structures of MC-1 and dCBP-1, we hypothesize that NOVA2 likely represents a protein highly sensitive to EP300 degradation in HAP-1 cells as opposed to an off-target. These findings define the proteome-wide selectivity of a paralogue-specific EP300 degrader.

**Figure 4.**
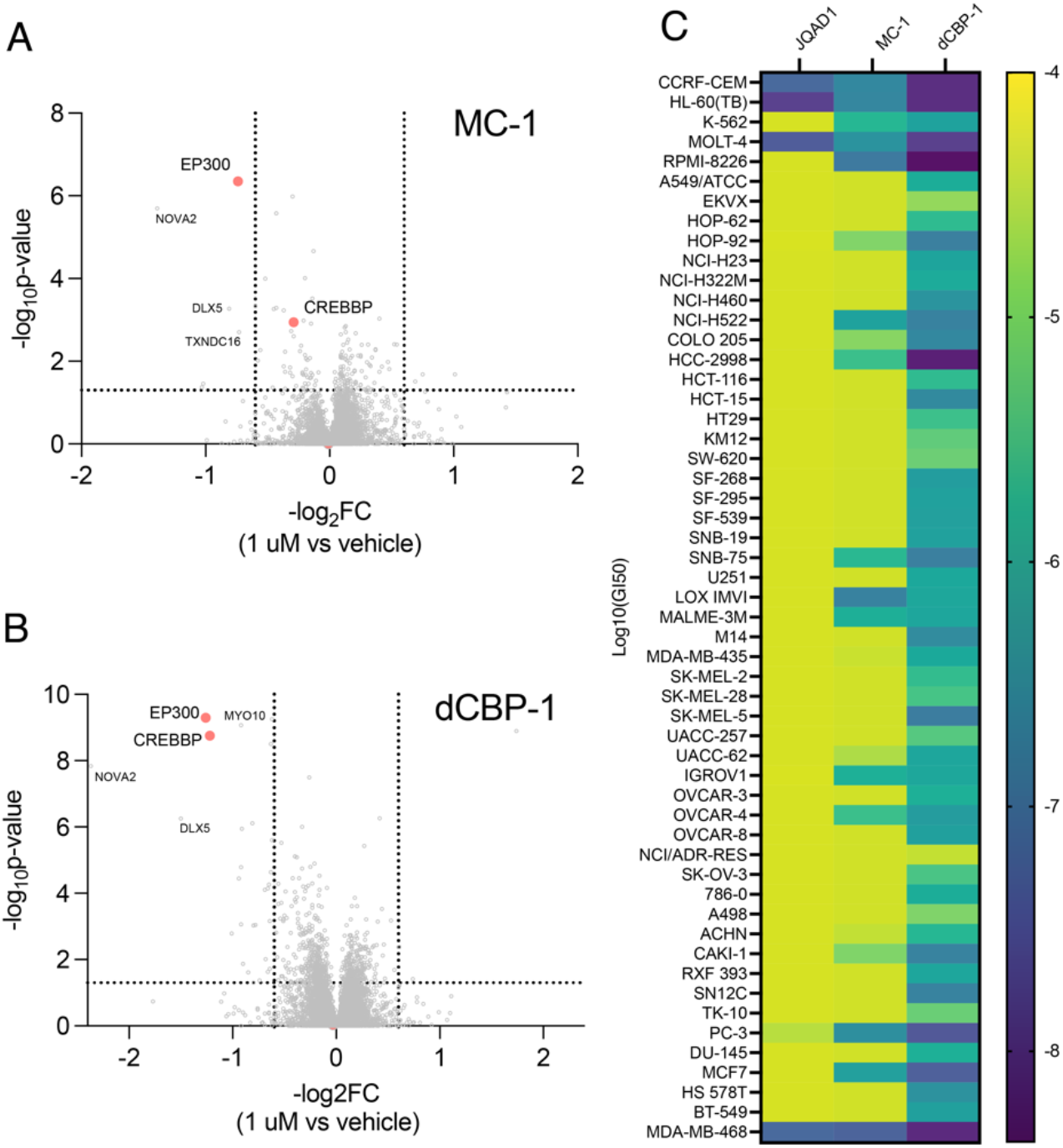
(A) Proteome-wide assessment of MC-1 treatment (1 μM, 6 h) in HAP1 cells with analysis of unique EP300/CREBBP peptides (n = 3 biological replicates). (B) Proteome-wide assessment of dCBP-1 treatment (1 μM, 6 h) in HAP1 cells with analysis of unique EP300/CREBBP peptides (n = 3 biological replicates). (C) Comparative analysis of half-maximal growth inhibition (GI_50_) values for EP300 degrader molecules across the NCI-60 cell line panel (72 hour incubation).

Next, MC1’s activity was tested in a broader context by examining its activity in the NCI60 cell line screen (Fig. 4b). This analysis used a recently redesigned, miniaturized version of the screen which employs a 384-well plate format, ATP-based viability readout, and 72 hour treatment regimen.^25^ To benchmark MC-1’s activity we also tested the previously reported dual EP300/CREBBP degrader dCBP-1 as well as the EP300-specific degrader JQAD-1. All three compounds were tested across five concentrations in order to calculate half-maximal growth inhibition (GI_50_). All three molecules were active against HL-60, CCRF-CEM, and MOLT-4 cells. This is consistent with the known dependence of hemopoietic malignancies on acetylation-dependent transcription.^1^ In terms of broader trends, MC-1 appeared to potently inhibit a unique subset of cell lines relative to JQAD-1 and dCBP-1 which were narrowly and broadly active, respectively. Quantitative dose-response profiling confirmed that MC-1 inhibited cell proliferation in MCF-7 cells, while HAP-1 cells were relatively unaffected (Fig. S8). Understanding that growth inhibition could be the result of either degradation or catalytic inhibition, we examined the dose-dependent effects of MC-1 on EP300, CREBBP, and H3K18Ac in these models. In HAP-1 cells, EP300 degradation occurs at low concentrations and appears to slightly precede complete loss of H3K18Ac caused by the dual inhibition of the EP300 and CREBBP HAT domains (Fig. 5a). MCF-7 cells display the opposite profile. Here, degradation occurs only to a limited extent at higher concentrations and appears to trail HAT inhibition (Fig. 5b). We hypothesize this may reflect different degradation kinetics in the two cell lines (Fig. 5c), potentially driven by increased drug efflux, reduced E3 ligase levels, or rapid rates of EP300 re-synthesis. Previous studies have mentioned the potential for specific degraders cause concentration-dependent catalytic inhibition,^12, 26^ although this has rarely been explored across multiple cell lines. Our results confirm the activity of paralogue-specific EP300 degraders across multiple cellular models and provide an approach for assessing dual inhibition versus paralogue-specific degradation.

**Figure 5.**
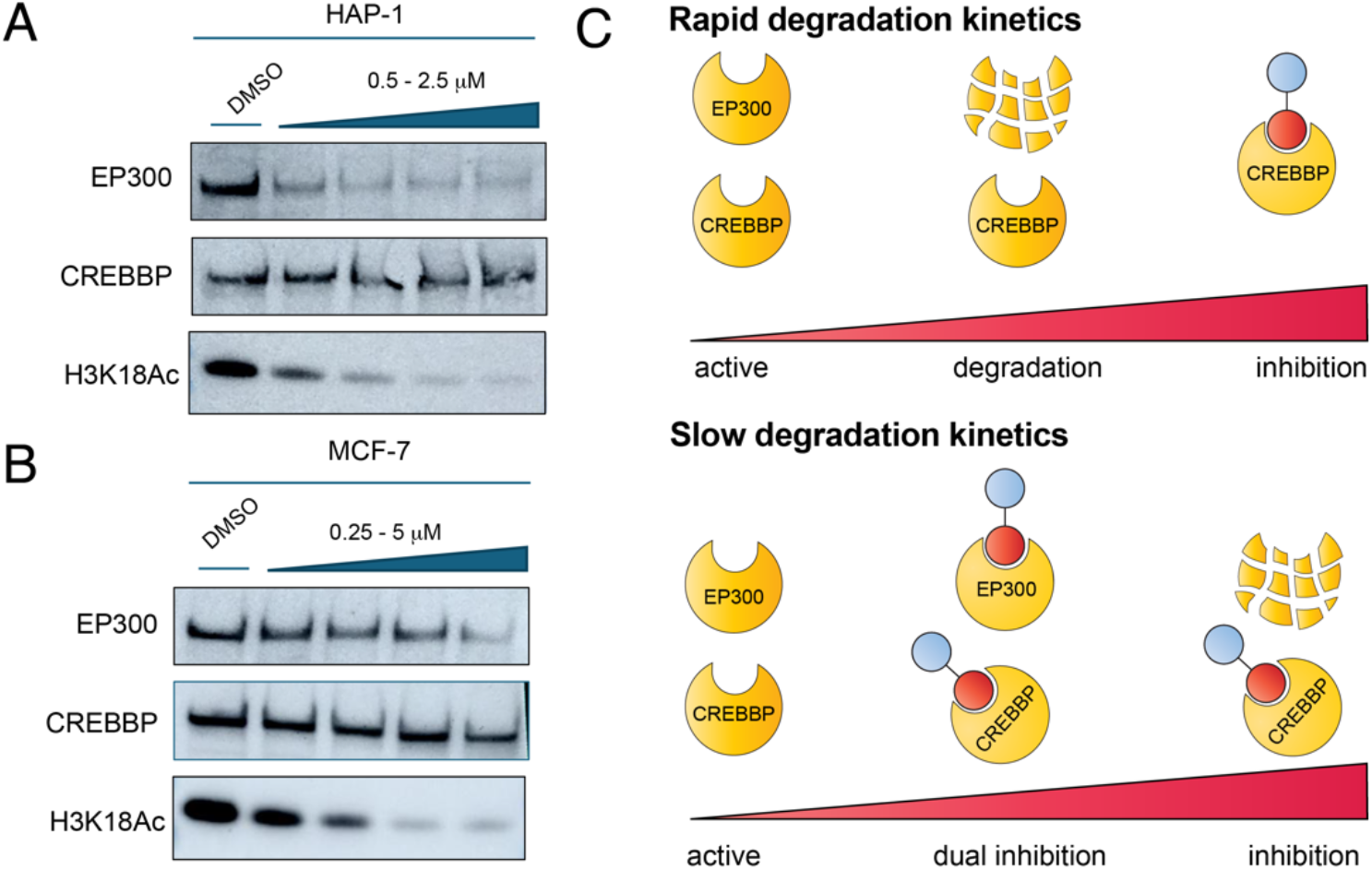
(A) Comparative effects of MC-1 on EP300 degradation and histone acetylation in HAP-1 cells. MC-1 was dosed at 0.25, 0.5, 1, and 2.5 μM for 6 h. (B) Comparative effects of MC-1 on EP300 degradation and histone acetylation in MCF-7 cells. MC-1 was dosed at 0.5, 1, 2.5, and 5 μM for 6 h (higher concentrations than in HAP-1 cells to observe clear degradation and inhibition). (C) Concentration and cell-dependent effects of HAT-based degraders. In cells where EP300 displays rapid degradation kinetics (HAP-1), MC-1 can significantly deplete EP300 while leaving CREBBP HAT activity intact. In cells where EP300 displays slower degradation kinetics (MCF-7), MC-1 can co-occupy and inhibit EP300 and CREBBP prior to EP300 degradation.

Here we report selective chemical degradation of the histone acetyltransferase EP300. Our studies complement significant previous efforts in the field but differentiate themselves by demonstrating for the first time that a potent HAT ligand in combination with VHL recruitment affords preferential degradation of EP300 over its closely related paralogue, CREBBP. Mechanistic studies indicate that selective degradation of EP300 by a bifunctional ligand cannot be predicted by measurement of target engagement (e.g. H3K18Ac) or ability to form a HAT-VHL-degrader ternary complex (e.g. NanoBiT). Instead, our results are most consistent with EP300’s i) slightly increased binding affinity for CPI-1612, ii) higher confidence predicted interactions with VHL, and iii) possession of two non-conserved lysine residues relative to CREBBP. Previous studies of p38 MAPK degraders have found ternary complex formation is necessary but not sufficient to explain degrader selectivity.^27^ Defining the downstream factors that govern selective EP300 degradation will be an important step for extending and improving this approach.

As currently constituted, we envision MC-1 will find utility studies aimed at differentiating the non-catalytic roles of EP300 and CREBBP, particularly in cell lines which are not greatly affected by loss of EP300/CREBBP HAT activity. In addition, it is known that ligands targeting the HAT domain of EP300/CREBBP are sensitive to cellular acetyl-CoA levels.^10, 28^ This suggests that the sensitivity of EP300 to MC-1 under different metabolic conditions may be a useful proxy for levels of cellular acetyl-CoA, an approach that could provide valuable insights into metabolic signaling.^29^

We also note limitations of our study and future directions of inquiry. MC-1 does not completely abolish cellular EP300. It caused a maximal degradation of ∼85% in our HAP-1 model and considerably less in MCF-7 cells. This, along with the observation that JQAD-1 was inactive in HAP-1 cells (Fig. S2), raises the question as to the degree to which cell line specificity may be a broader attribute of paralogue-specific EP300 degraders. For example, JQAD-1’s mechanism was recently validated in a study showing that CRBN KO confers resistance in several models,^30^ but this was observed in only 9/19 cell lines tested. MC-1’s ability to inhibit H3K18Ac in MCF-7 would argue against differential occupancy being the sole determinant of this variability. Other criteria that may influence degradation include cellular accumulation, E3 ligase activity, and the ability of P300 partner proteins^31^ to influence accessibility and correct orientation of the ubiqutinylation machinery. During preparation of this manuscript a CPI-1612-derived CRBN-based dual EP300/CREBBP degrader was reported.^21^ This ever-expanding toolbox of HAT domain-targeting degraders, in combination with chemical and genetic screens, should be helpful in identifying factors underlying cell-type specific degradation.

Our studies also emphasize the emerging spectrum of options for therapeutic EP300/CREBBP inhibition. In addition to the well-characterized HAT^10^ and BRD^32-34^ ligands, there are now multiple potent dual degraders based on BRD ligands.^13-16, 35^ JQAD-1 and MC-1 add to this collection by affording paralogue-specific degradation and – in the case of MC-1, and possibly JQAD-1 as well – dual HAT inhibition at high concentrations. In this latter scenario, MC-1 may be deployed as an augmented HAT inhibitor with an additional capability to degrade EP300. Interestingly, a recent study reported that dual EP300/CREBBP degraders do not cause weight loss in mice.^36^ This implies either that dual loss of EP300/CREBBP catalytic activity is more well-tolerated than expected, or that the unique pharmacology of bifunctional molecules may spare EP300/CREBBP in settings where it is essential. VHL-linked degraders have shown the capacity to mitigate thrombocytopenia caused by their parent inhibitor compounds in platelet models.^37^ Whether MC-1 or improved derivatives will show similarly altered safety or efficacy is currently unknown. These studies are underway and will be reported in due course.

## Acknowledgements

The authors thank Dr. Francis O’Reilly (NCI) for helpful discussions. We are grateful to Prof. Jun Qi (DFCI) for supplying JQAD-1. This work was supported by the Intramural Research Programs of the National Cancer Institute, Center for Cancer Research ZIA BC011488. This work utilized the computational resources of the NIH HPC Biowulf cluster (http://hpc.nih.gov).

## Supplementary Figures & Tables

**Table S1.**
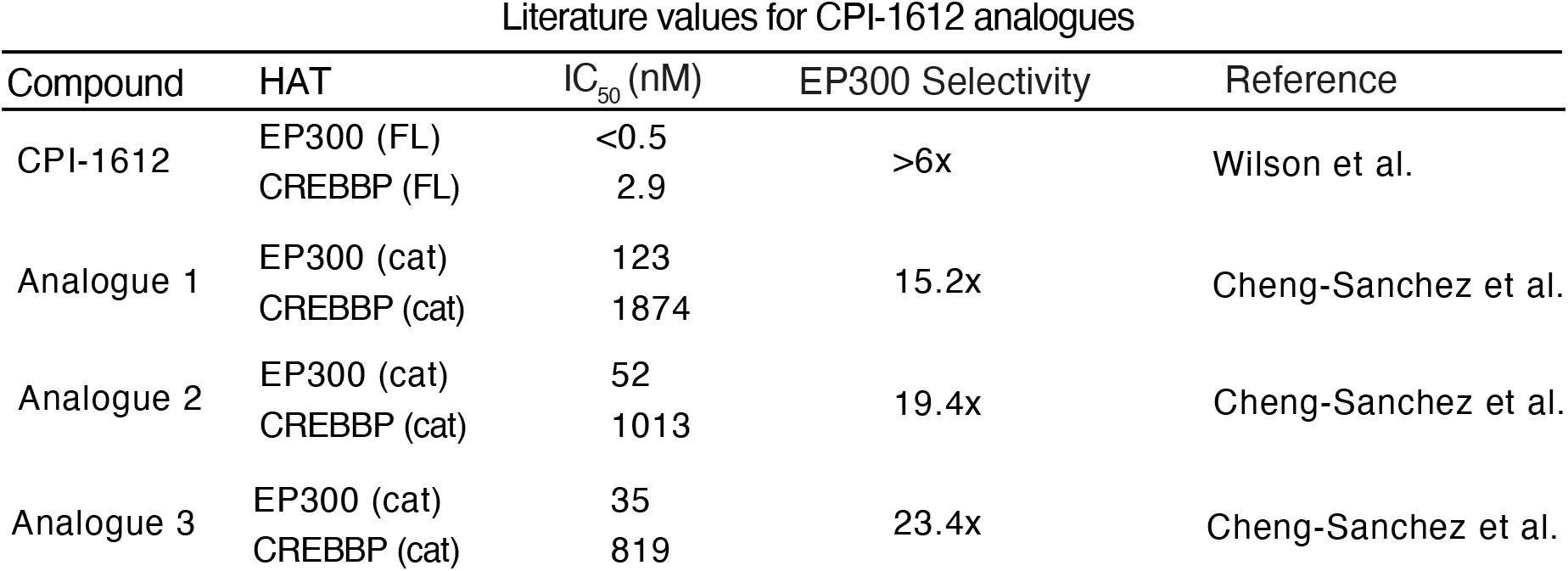
Literature values comparing biochemical inhibition of EP300 and CREBBP by small molecule using the CPI-1612 scaffold.

**Figure S1.**
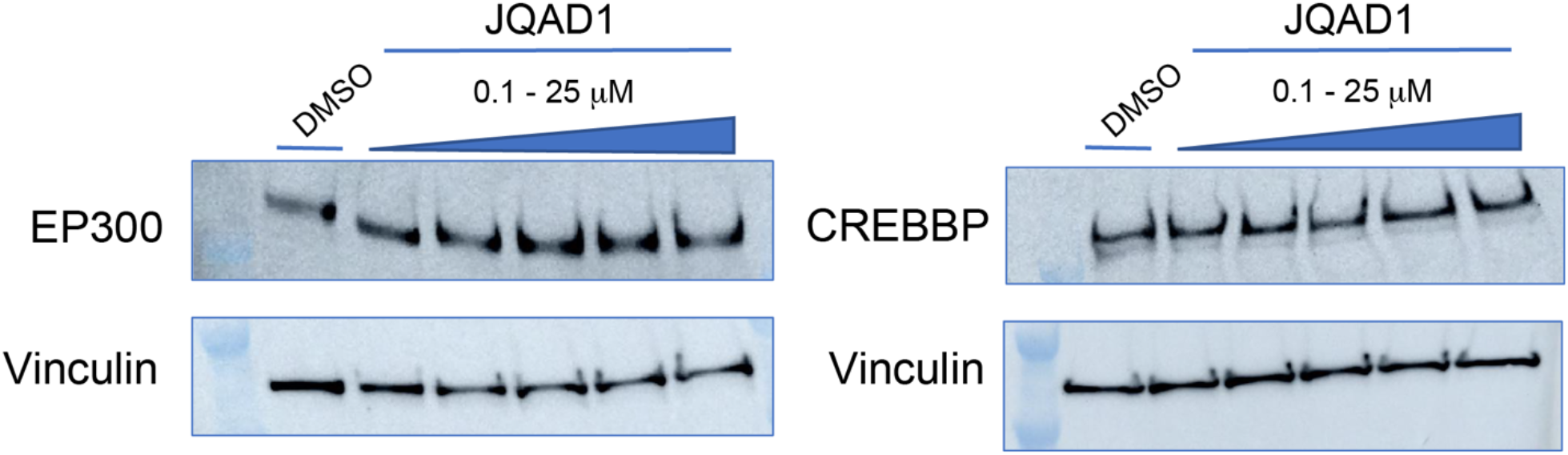
JQAD-1 does not appear to degrade EP300 or CREBBP in HAP-1 cells after a 6 h incubation. Concentrations = 0.1, 1, 5, 10, 25 µM.

**Figure S2.**
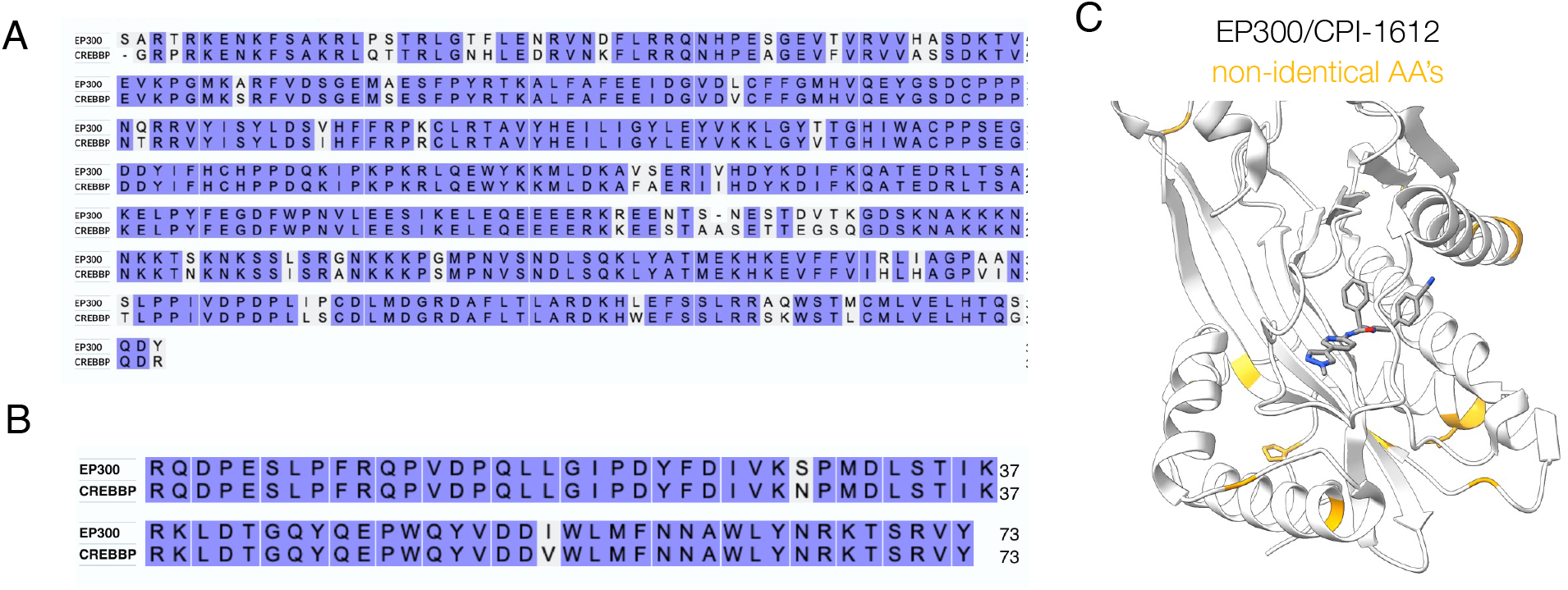
(a) Sequence alignment of EP300 and CREBBP HAT domains. (b) Sequence alignment of EP300 and CREBBP bromodomains. (c) Crystal structure of EP300 HAT domain bound to CPI-1612 (PDB: 6V8N). Residues in yellow represent amino acids that are not conserved between EP300 and CREBBP.

**Figure S3.**
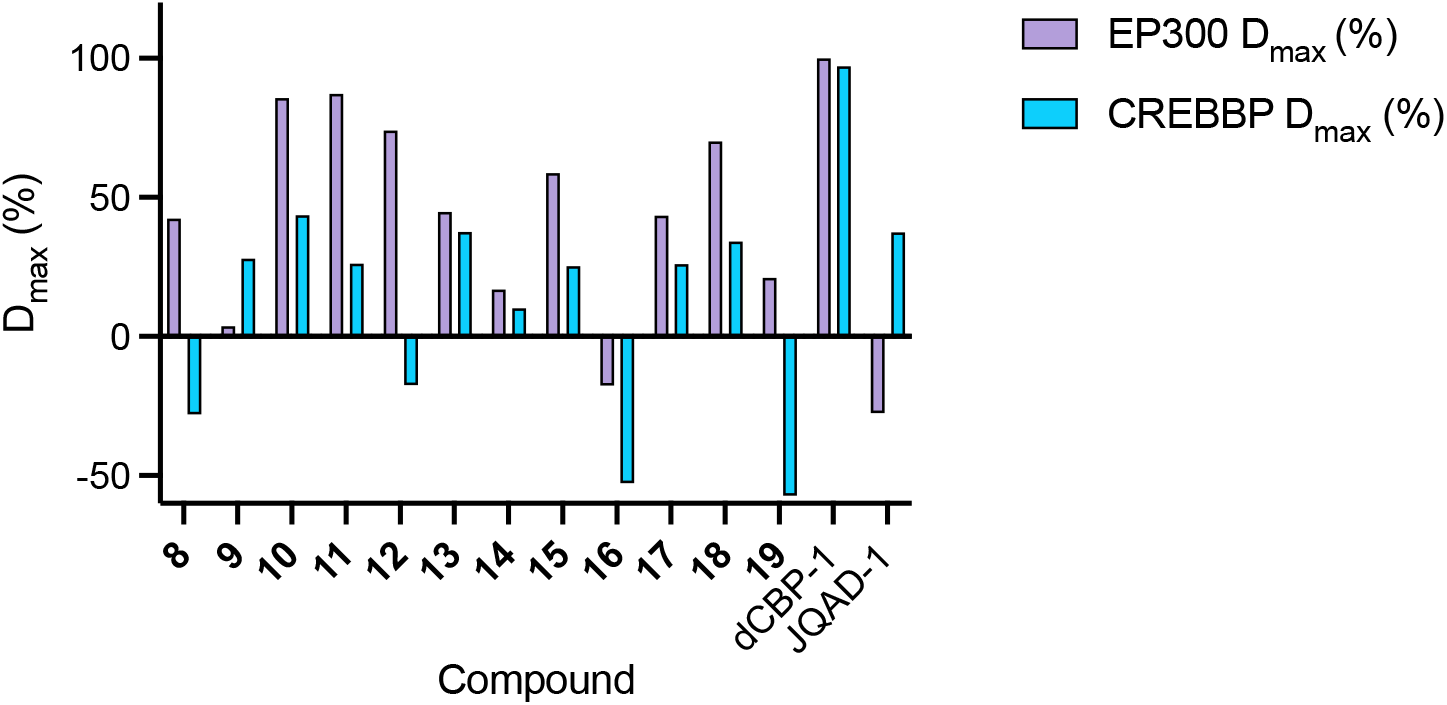
D_max_of EP300 and CREBBP approximated by gel densitometry from primary degrader screen Western blots in Fig. 2B or Fig. S1 (n = 1). Quantification of **8**-**19** were from ∼24 h treatments with concentrations of 0.5, 1, 5, and 10 µM, dCBP-1 was treated for 24 h with concentrations of 0.001, 0.01, 0.1, 1, and 10 µM, and JQAD-1 was treated for 6 h with concentrations of 0.1, 1, 5, 10, and 25 µM. Among **8**-**19**, all compounds except **9** show stronger degradation of EP300 than CREBBP, with compounds **10** and **11** showing most potent EP300 degradation.

**Figure S4.**
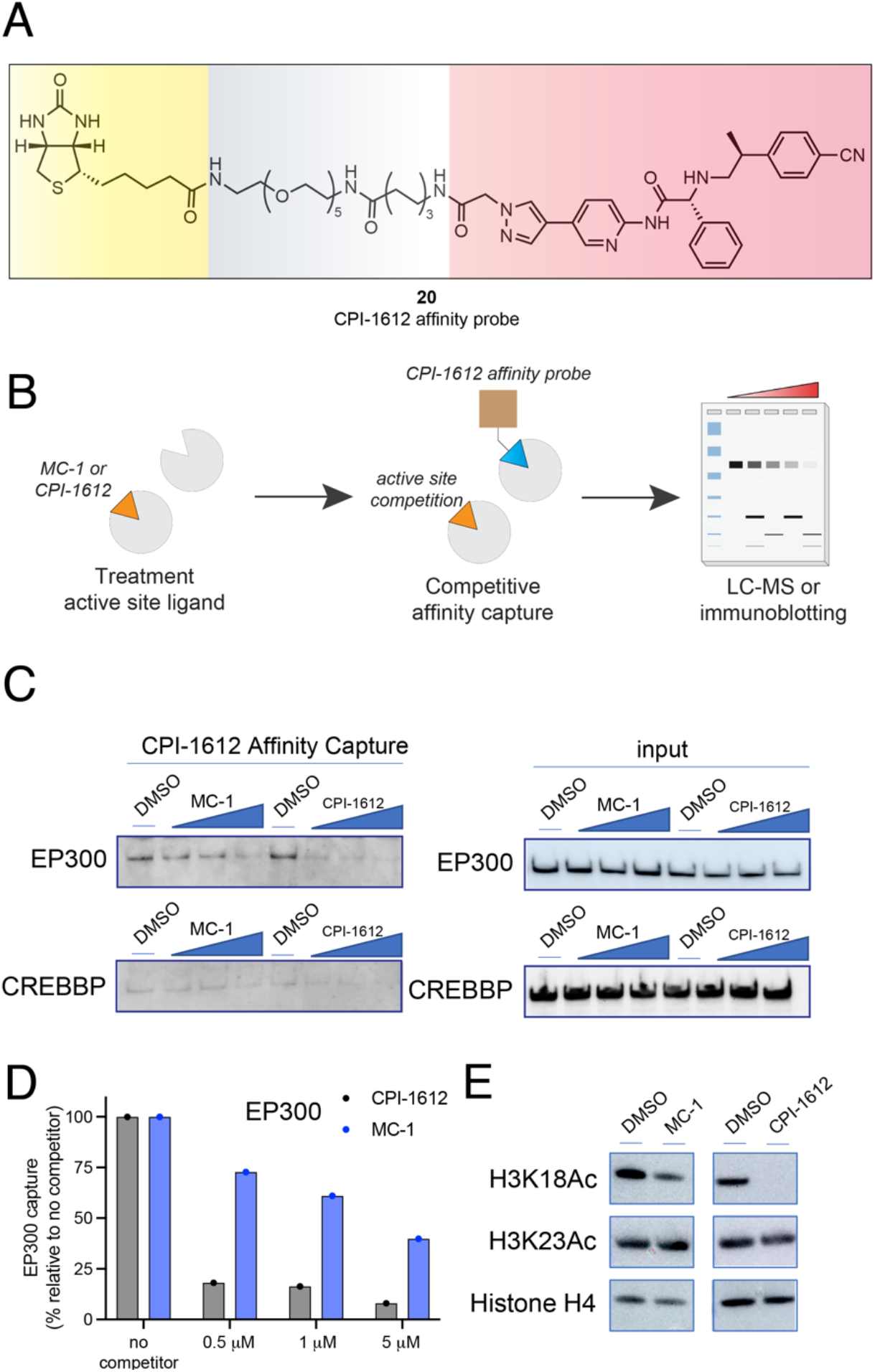
(A) Structure of CPI-1612 affinity probe. (B) Evaluating the relative affinity of CPI-1612 and MC-1 for endogenous EP300/CREBBP by competitive affinity capture. Nuclear extracts are pre-incubated with small molecule prior to addition of CPI-1612 affinity probe which is CPI-1612 coated streptavidin beads. Following affinity capture, proteins are eluted and assessed for levels of EP300 and CREBBP. (C) Western blotting data evaluating EP300 and CREBBP affinity capture in the presence of DMSO, increasing MC-1, or increasing CPI-1612. Concentrations of MC-1 or CPI-1612: 0.5, 1, 5 μM. EP300 and CREBBP levels in input samples for affinity capture experiment are shown on the right. (D) Quantified affinity capture of EP300 in the presence of increasing CPI-1612 (grey) or MC-1 (blue). Data was from Fig.S3C and quantified by ImageJ. (E) Comparative effects of CPI-1612 and MC-1 on EP300/CREBBP-catalyzed histone H3K18 acetylation in HAP1 cells. Both compounds were dosed at 250 nM for 6 h.

**Figure S5.**
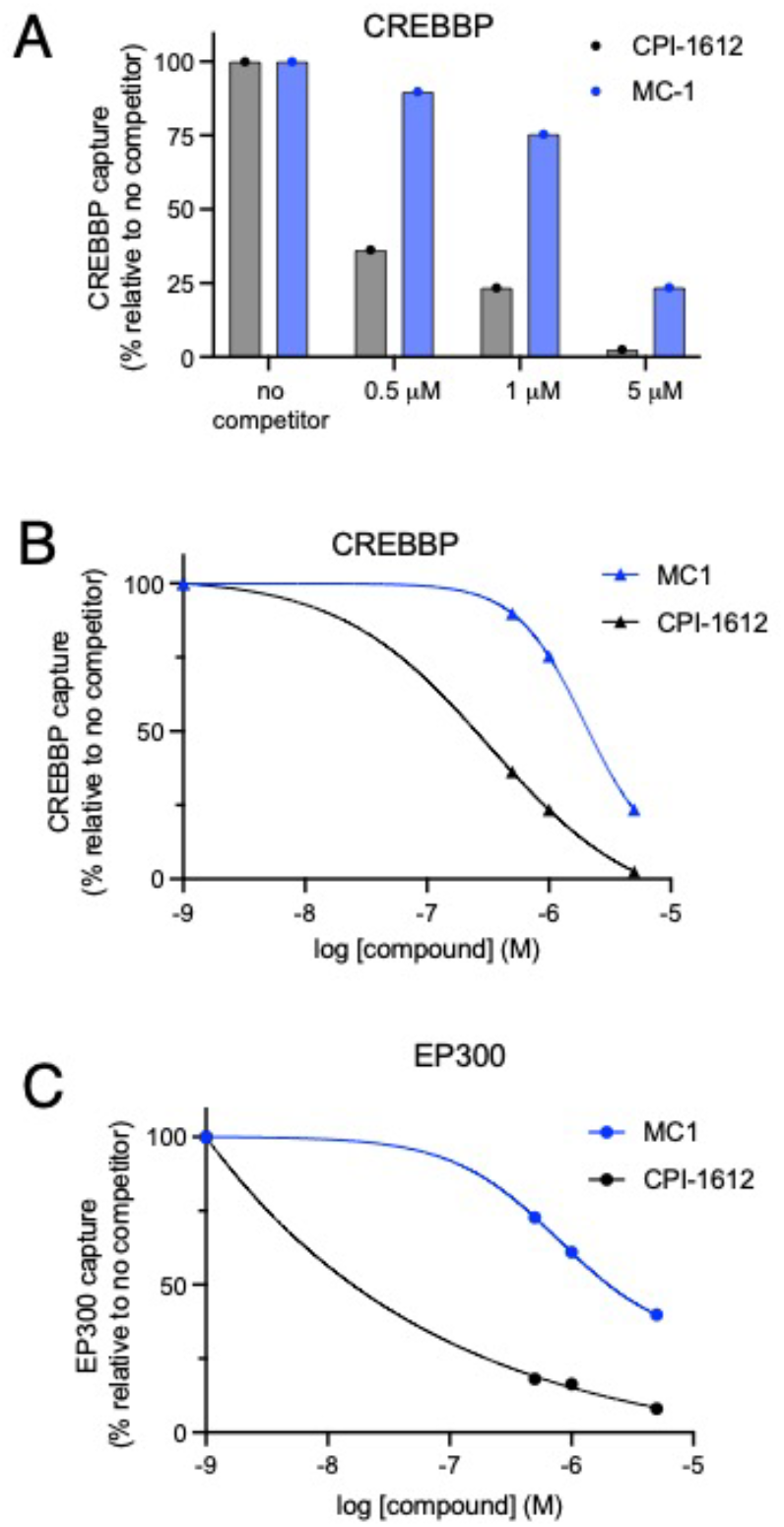
(A) Quantified affinity capture of CREBBP in the presence of increasing CPI-1612 (grey) or MC-1 (blue). Data was from Fig.S3C and quantified by ImageJ. (B) Quantified affinity capture of CREBBP in the presence of increasing CPI-1612 (grey) or MC-1 (blue) fit using non-linear regression model. (C) Quantified affinity capture of EP300 in the presence of increasing CPI-1612 (grey) or MC-1 (blue) fit using non-linear regression model. Concentrations of MC-1 or CPI-1612: 0, 0.5, 1, 5 mM. Quantified data was from Fig.S3C and measured by ImageJ then processed in GraphPad Prism 10 by using non-linear regression model.(B,C). CPI-1612 and MC-1 appear to more potently antagonize EP300 capture (relative to CREBBP) in this experiment.

**Figure S6.**
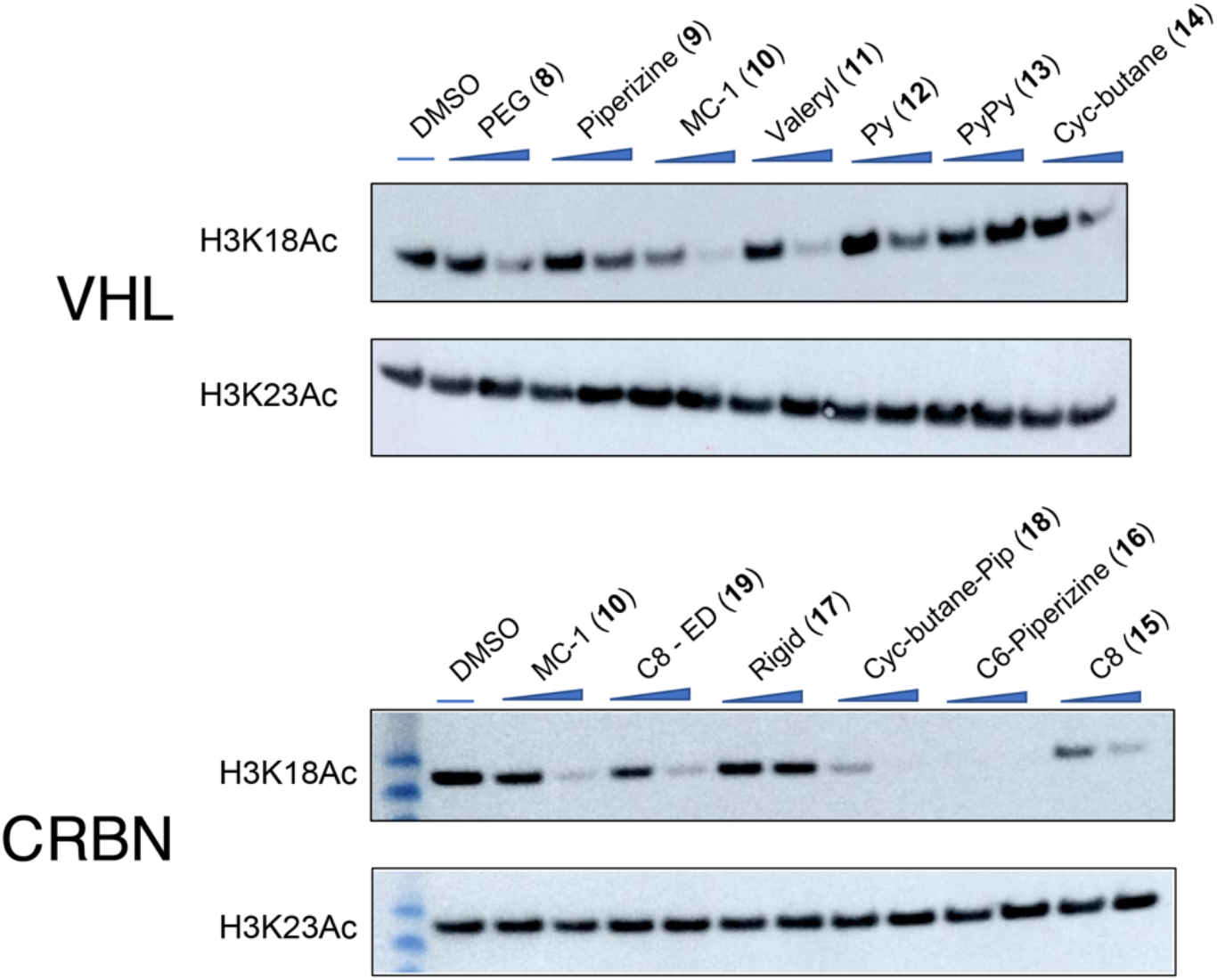
Evaluating the ability of CPI-1612-based degraders using VHL (top) and CRBN (bottom) E3 ligase recruiters to inhibit EP300/CREBBP-catalyzed histone H3K18Ac in HAP1 cells. Cells were treated for 6 h with either 250 nM or 2500 nM of the indicated compound. Inhibition of H3K18Ac provides a proxy for inhibitor potency and uptake. A lack of inhibition indicates a problem with inhibitory potency, uptake, or both.

**Figure S7.**
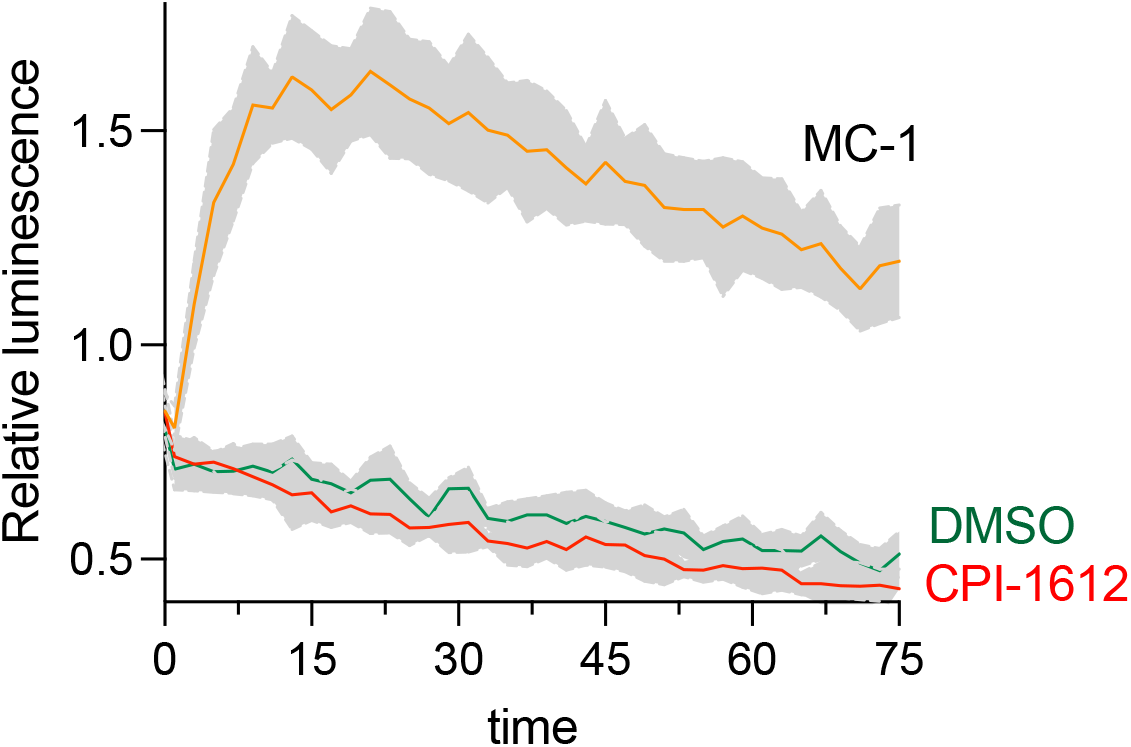
Measurement of CREBBP-VHL binding via NanoBiT assay. Averages of 4 replicates are plotted, shaded areas represent standard deviation of mean.

**Figure S8.**
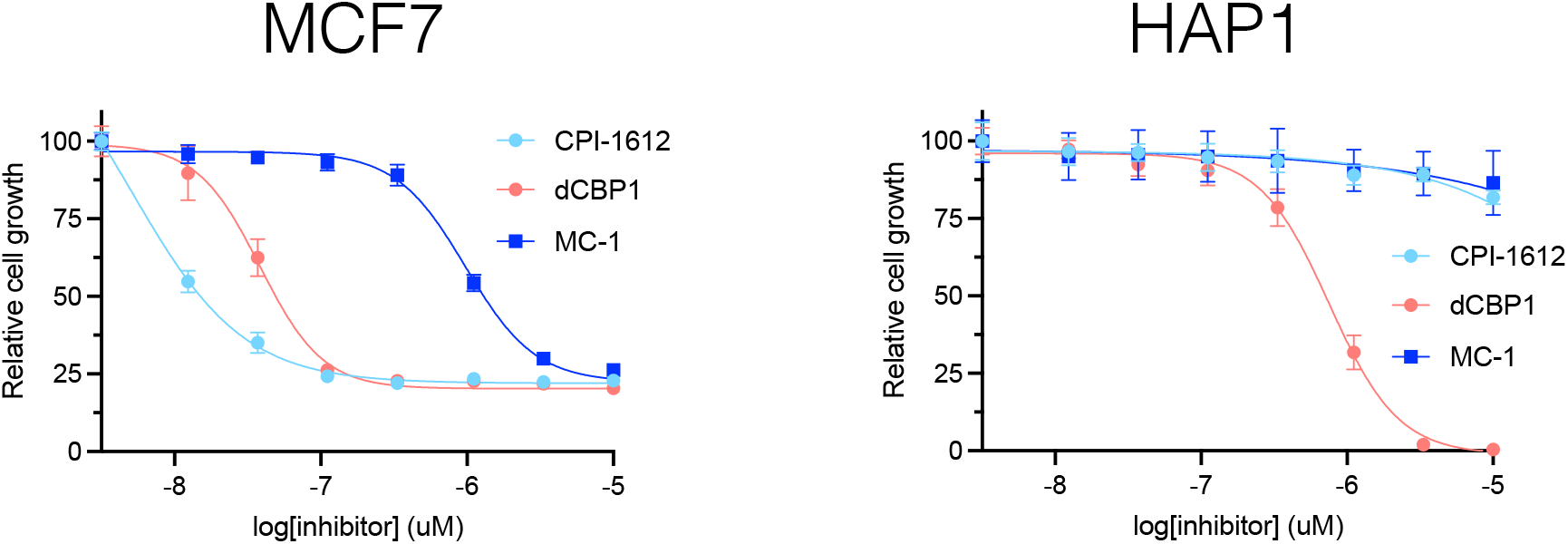
Relative cytotoxicity of CPI-1612, dCBP1, and MC-1 in HAP1 and MCF7 cells. Cell viability/growth was measured by Cell Titer Glo assay after 96 h incubation. Relative cell viability/growth was normalized to DMSO vehicle controls. n = 4 biological replicates.

## Notes

### Competing Interest Statement

The authors have declared no competing interest.

