## Supporting Information for "Paralogue-selective degradation of the lysine acetyltransferase EP300"

**Table of Contents for Supporting Information**

| | Page |
|---|---|
| Supplementary figures & tables | S2 |
| General materials and methods | S9 |
| Synthesis of degrader molecules | S11 |
| Primary degrader screen | S15 |
| Competitive affinity pulldown protocol | S15 |
| Histone modification analyses | S15 |
| Mechanistic analyses of degrader molecules | S16 |
| LC-MS/MS analyses of specificity | S18 |
| Full Western blot images | S21 |
| References | S35 |

### Supplementary Figures & Tables

Literature values for CPI-1612 analogues

| Compound | HAT | $IC_{50}$ (nM) | EP300 Selectivity | Reference |
|---|---|---|---|---|
| CPI-1612 | EP300 (FL) | <0.5 | >6x | Wilson et al. |
| | CREBBP (FL) | 2.9 | | |
| Analogue 1 | EP300 (cat) | 123 | 15.2x | Cheng-Sanchez et al. |
| | CREBBP (cat) | 1874 | | |
| Analogue 2 | EP300 (cat) | 52 | 19.4x | Cheng-Sanchez et al. |
| | CREBBP (cat) | 1013 | | |
| Analogue 3 | EP300 (cat) | 35 | 23.4x | Cheng-Sanchez et al. |
| | CREBBP (cat) | 819 | | |

**Table S1.** Literature values comparing biochemical inhibition of EP300 and CREBBP by small molecule using the CPI-1612 scaffold.[1, 2]

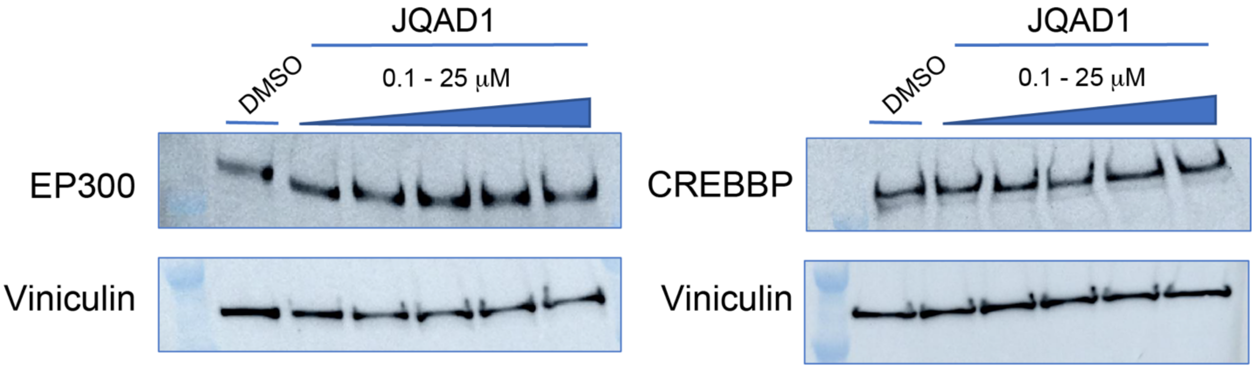


**Figure S1**. JQAD-1 does not appear to degrade EP300 or CREBBP in HAP-1 cells after a 6 h incubation. Concentrations = 0.1, 1, 5, 10, 25 μM.

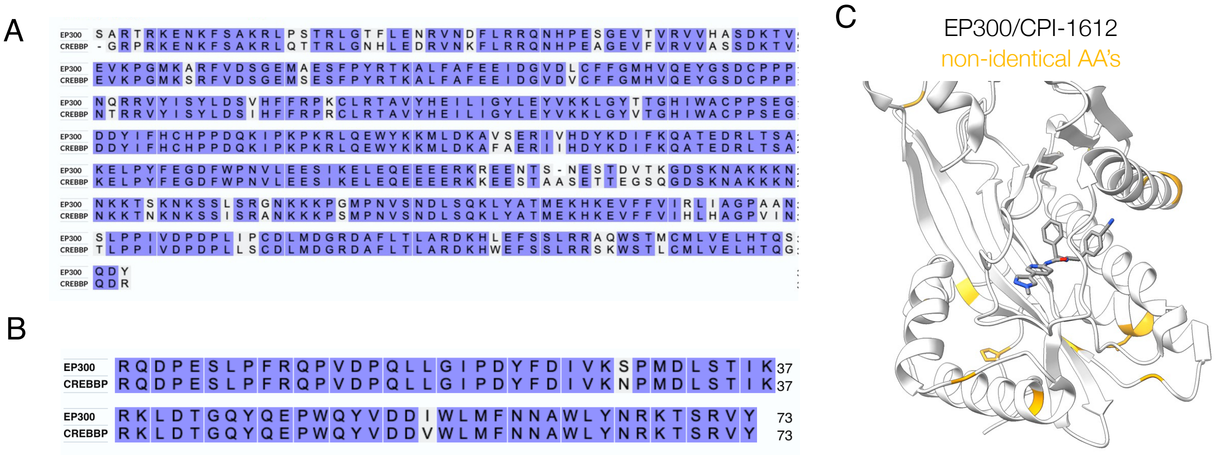


**Figure S2.** (a) Sequence alignment of EP300 and CREBBP HAT domains. (b) Sequence alignment of EP300 and CREBBP bromodomains. (c) Crystal structure of EP300 HAT domain bound to CPI-1612 (PDB: 6V8N). Residues in yellow represent amino acids that are not conserved between EP300 and CREBBP.

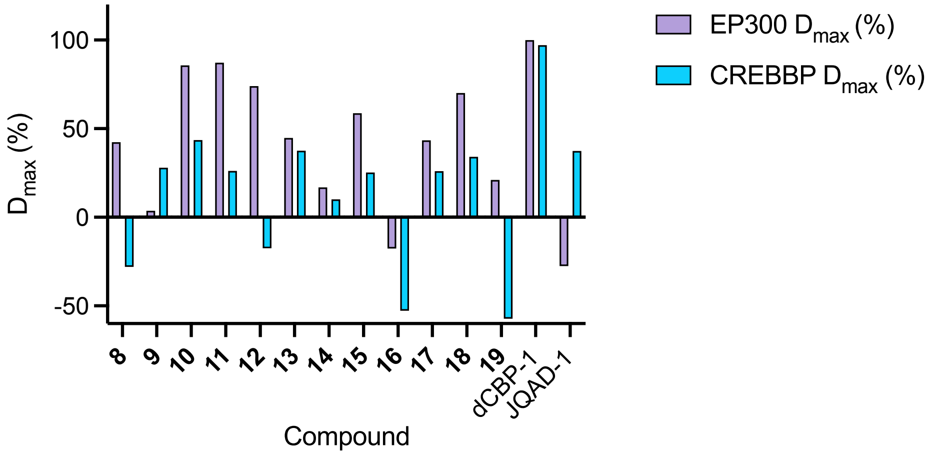


**Figure S3.** $D_{max}$ of EP300 and CREBBP approximated by gel densitometry from primary degrader screen Western blots in Fig. 2B or Fig. S1 (n = 1). Quantification of **8-19** were from ~24 h treatments with concentrations of 0.5, 1, 5, and 10 μM, dCBP-1 was treated for 24 h with concentrations of 0.001, 0.01, 0.1, 1, and 10 μM, and JQAD-1 was treated for 6 h with concentrations of 0.1, 1, 5, 10, and 25 μM. Among **8-19**, all compounds except **9** show stronger degradation of EP300 than CREBBP, with compounds **10** and **11** showing most potent EP300 degradation.

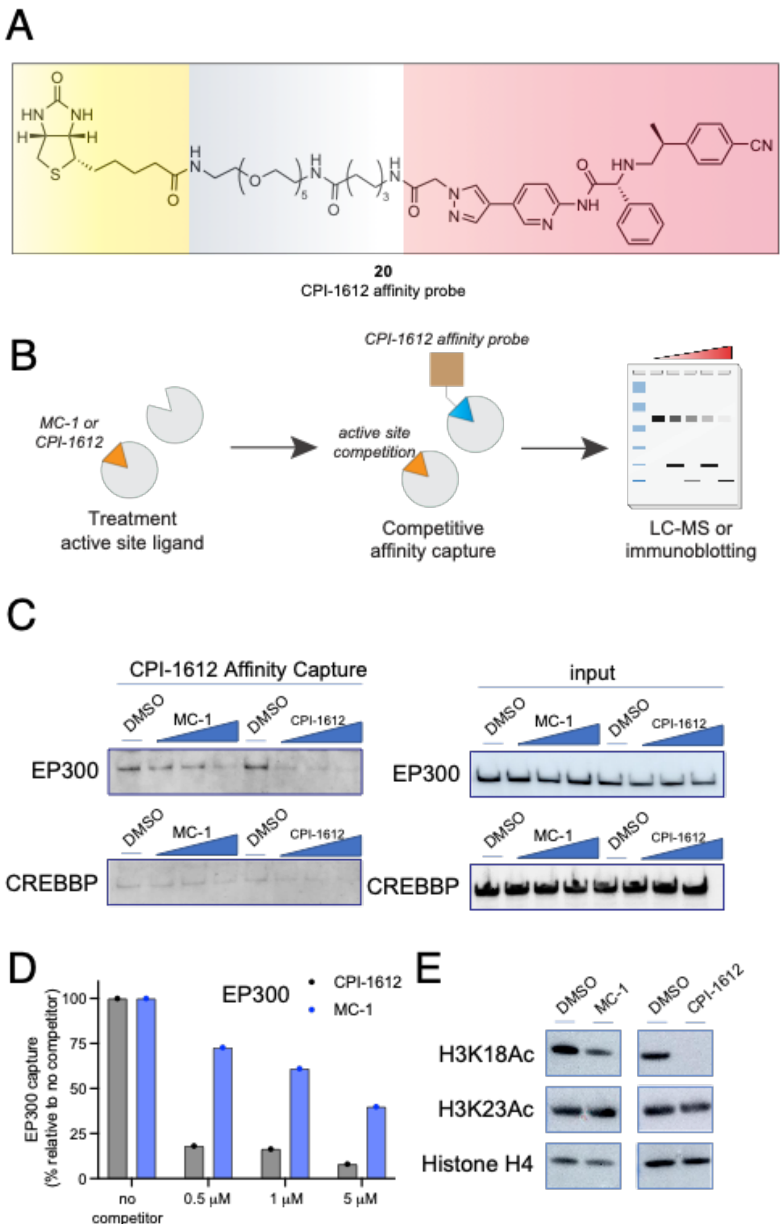


**Figure S4.** (A) Structure of CPI-1612 affinity probe. (B) Evaluating the relative affinity of CPI-1612 and MC-1 for endogenous EP300/CREBBP by competitive affinity capture. Nuclear extracts are pre-incubated with small molecule prior to addition of CPI-1612 affinity probe which is CPI-1612 coated streptavidin beads. Following affinity capture, proteins are eluted and assessed for levels of EP300 and CREBBP. (C) Western blotting data evaluating EP300 and CREBBP affinity capture in the presence of DMSO, increasing MC-1, or increasing CPI-1612. Concentrations of MC-1 or CPI-1612: 0.5, 1, 5 $\mu$M. EP300 and CREBBP levels in input samples for affinity capture experiment are shown on the right. (D) Quantified affinity capture of EP300 in the presence of increasing CPI-1612 (grey) or MC-1 (blue). Data was from Fig.S3C and quantified by ImageJ. (E) Comparative effects of CPI-1612 and MC-1 on EP300/CREBBP-catalyzed histone H3K18 acetylation in HAP1 cells. Both compounds were dosed at 250 nM for 6 h.

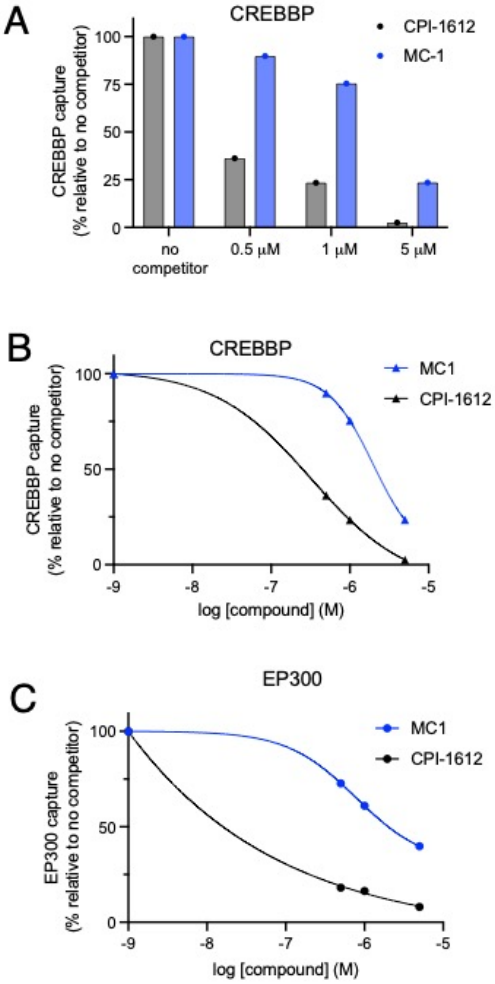


**Figure S5.** (A) Quantified affinity capture of CREBBP in the presence of increasing CPI-1612 (grey) or MC-1 (blue). Data was from Fig.S3C and quantified by ImageJ. (B) Quantified affinity capture of CREBBP in the presence of increasing CPI-1612 (grey) or MC-1 (blue) fit using non-linear regression model. (C) Quantified affinity capture of EP300 in the presence of increasing CPI-1612 (grey) or MC-1 (blue) fit using non-linear regression model. Concentrations of MC-1 or CPI-1612: 0, 0.5, 1, 5 mM. Quantified data was from Fig.S3C and measured by ImageJ then processed in GraphPad Prism 10 by using non-linear regression model.(B,C). CPI-1612 and MC-1 appear to more potently antagonize EP300 capture (relative to CREBBP) in this experiment.

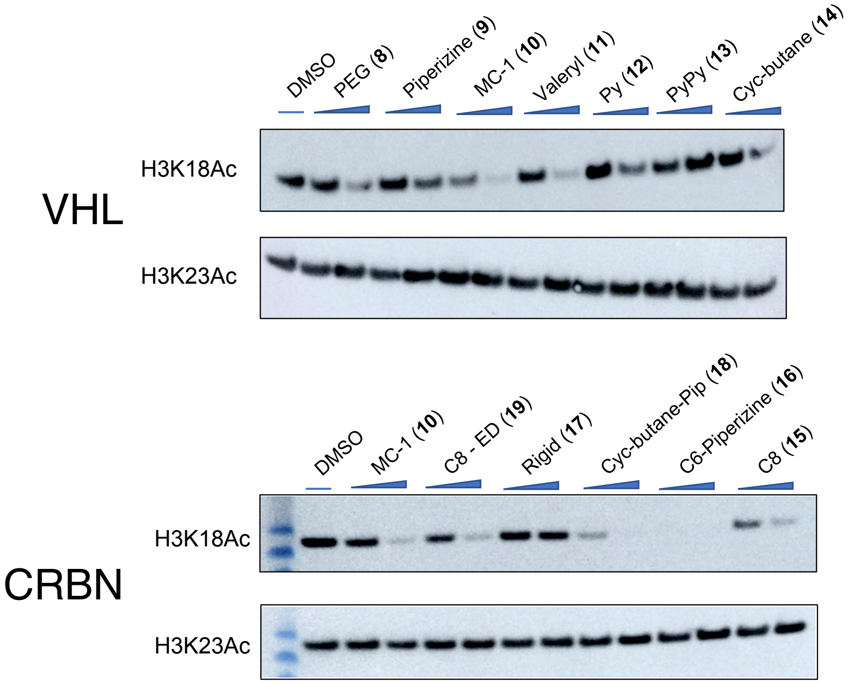


**Figure S6.** Evaluating the ability of CPI-1612-based degraders using VHL (top) and CRBN (bottom) E3 ligase recruiters to inhibit EP300/CREBBP-catalyzed histone H3K18Ac in HAP1 cells. Cells were treated for 6 h with either 250 nM or 2500 nM of the indicated compound. Inhibition of H3K18Ac provides a proxy for inhibitor potency and uptake. A lack of inhibition indicates a problem with inhibitory potency, uptake, or both.

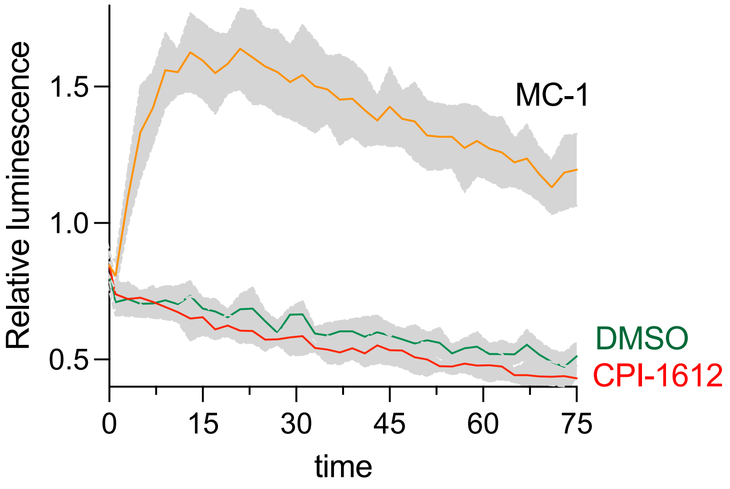


**Figure S7.** Measurement of CREBBP-VHL binding via NanoBiT assay. Averages of 4 replicates are plotted, shaded areas represent standard deviation of mean.

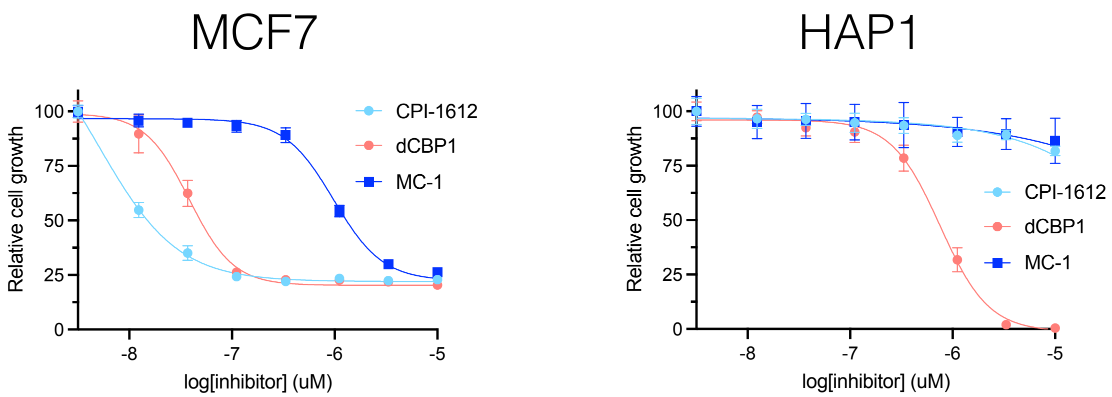


**Figure S8**. Relative cytotoxicity of CPI-1612, dCBP1, and MC-1 in HAP1 and MCF7 cells. Cell viability/growth was measured by Cell Titer Glo assay after 96 h incubation. Relative cell viability/growth was normalized to DMSO vehicle controls. n = 4 biological replicates.

### General materials and methods

Chemicals were purchased from Sigma Aldrich, Acros Organics, Alfa Aesar, Oakwood Chemical, Ambeed, Inc. or Chem Impex International. All nonaqueous reactions were carried out using flame- or oven-dried glassware under an atmosphere of dry argon or nitrogen. Tetrahydrofuran (THF), dichloromethane (DCM), *N,N*-dimethylformamide (DMF), and methanol (MeOH) were purified via filtration through two columns of activated basic alumina under an atmosphere of Argon using a solvent purification system from Pure Process Technology. Other commercial reagents were used as received unless otherwise noted. $^{1}$H Nuclear magnetic resonance ($^{1}$H NMR) and $^{13}$C NMR spectra were acquired on a Varian 400MHz instrument (400 MHz for $^{1}$H and 100 MHz for $^{13}$C). For $^{1}$H and $^{13}$C, chemical shifts (δ) are reported in ppm referenced to $CDCl_3$ (7.26 ppm for $^{1}$H and 77.2 ppm for $^{13}$C), $CD_3OD$ (3.31 ppm for $^{1}$H, 49.0 ppm for $^{13}$C), or dimethyl sulfoxide (DMSO)-$d_6$ (2.50 ppm for $^{1}$H, 39.5 ppm for $^{13}$C). LC/MS data were secured from Agilent 1200 LC/MS spectrometer. The HRMS analysis was achieved on a Waters Xevo G2-XS using ESI ionization. Thin layer chromatography (TLC) was performed using EMD aluminum-backed (0.20 mm) silica plates (60 F-254), and flash chromatography used ICN silica gel (200−400 mesh). dCBP-1 (HY-134582) and CPI-1612 (HY-136285) were purchased from MedChemExpress. JQAD-1 used in the NCI-60 screen was purchased from Tocris (76-825). JQAD-1 used in Western blotting experiments was a kind gift of Prof. Jun Qi (Dana-Farber Cancer Institute).

Sequence homology and identity were calculated by aligning the Uniprot-defined EP300 and CREBBP bromodomains (EP300 1067-1139, CREBBP 1103-1175) and HAT domains (EP300 1287-1663, CREBBP 1323-1700) using the ClustalOmega server. Aligned sequence outputs from ClustalOmega were used in FASTA format as an input for calculation of identity and similarity using Sequence Manipulation Suite (https://www.bioinformatics.org/sms2/ident_sim.html). Alphafold Multimer predictions were performed using the protein sequence of the following Uniprot entries: catalytic core of EP300 (Q09472, 1048-1665), catalytic core of CREBBP (Q92793, 1084-1702), CRBN (Q96SW2), and VHL (P40337). Analyses were carried out using the Collabfold implementation on the NIH's Biowulf high-performance computing sytem using the default settings. The complexes were predicted without template information. MSA options were set as follows: MMseqs. 2 (UniRef + Environmental) was chosen as MSA mode and unpaired + paired as pair mode, and the model type was set to auto. Further documentation can be found at: https://hpc.nih.gov/apps/colabfold.html.

HAP-1, MCF-7, and HEK-293T cells were cultured at 37 °C under 5% $CO_2$, and all were passaged every 2–3 days at 70–90% confluency. HAP-1 cells were grown in IMDM (Thermo 12440053) with 10% FBS (Avantor Seradigm 97068-085) and 1% penicillin-streptomycin (Thermo 15140122), MCF-7 cells were grown in EMEM (Quality Biological 112-016-101) with 10% FBS, 2 mM L-glutamine (Quality Biological 118-084-721), 0.01 mg/mL human recombinant insulin (MilliporeSigma 91077C), and 1% penicillin-streptomycin, and HEK-293T cells were grown in

DMEM (Quality Biological 112-013-101) with 10% FBS, 2 mM L-glutamine and 1% penicillin-streptomycin.

Cells were lysed for Western blotting of EP300/CREBBP in 1x RIPA buffer (Abcam ab156034) supplemented with 1x protease inhibitor cocktail (Cell Signaling Technology 5871). Precision Red Protein Assay (Cytoskeleton ADV02) was used for protein quantification prior to Western blots. SDS-PAGE was performed using NuPAGE 3 to 8% Tris-Acetate gels (Invitrogen EA03752BOX and EA03755BOX) for EP300/CREBBP gels, or 4-12% Bis-Tris NuPAGE gels (Invitrogen NP0322 and NP0323) for histone gels. Gels were run with XCell SureLock Mini-Cells (Invitrogen EI0002) with Tris-Acetate SDS running buffer (Invitrogen LA0041) for Tris-Acetate gels, or MES running buffer (Invitrogen NP0002) for Bis-Tris. Precision Plus protein ladder (Bio-Rad 1610374) or Spectra Multicolor Broad Range protein ladder (Thermo 26634) were used for Tris-Acetate gels, and BenchMark Pre-stained protein ladder was used for Bis-Tris gels (Invitrogen 10748010).

For Western blotting, Tris-Acetate gels were transferred to 0.45 μM pore nitrocellulose membranes (Novex, Life Technologies # LC2001) by wet transfer electroblotting at 30 volts for 1 h using a XCell II Blot Module (Invitrogen EI9051) and NuPAGE transfer buffer (Invitrogen NP00061) following the manufacturer's protocols for Tris-Acetate gels. Bis-Tris gels were transferred using the iBlot dry blotting system (Invitrogen IB1001) and nitrocellulose transfer stacks (Invitrogen IB301001) at 20 volts for 8 min. Total protein on Western blots was visualized with Ponceau staining followed by washing of membranes with 5% acetic acid in $ddH_2O$. To visualize histone acetylation, membranes were blocked using StartingBlock (PBS) Blocking Buffer (Thermo Scientific 37538) for 20 min at room temperature, and then incubated overnight at 4 °C in a solution containing the primary antibody at the indicated dilution in StartingBlock Blocking Buffer. EP300 (Santa Cruz SC-584) and CREBBP (Cell Signaling Technologies 7389S) antibodies were used at 1:1,000 dilutions, while H3K18Ac (07-354) and H3K23Ac (07-355) antibodies were purchased from MilliporeSigma and used at 1:10,000 dilutions. Separate blots were run for EP300 versus CREBBP and for H318Ac versus H3K23Ac due to their shared molecular weights and secondary antibodies, while Vinculin-HRP (Cell Signaling 18799) at a 1:1,000 dilution was probed on both blots without stripping after imaging for EP300 or CREBBP. Since Vinculin results were similar on both blots, only one may be shown in figures. Membranes were washed the next day with 1x TBST at least 3 times and incubated for 1 h with anti-rabbit IgG HRP-linked antibody (Cell Signaling Technology 7074S) diluted to 1:1,000 in 1x TBST + 5% non-fat dry milk. Membranes were washed at least 3 times with 1x TBST, prior to imaging with Lumiglo (Cell Signaling Technology #7003) or SuperSignal ELISA Femto Substrate (Thermo Scientific 37074) according to manufacturer's protocols. Colorimetric and chemiluminescent signals were detected using an Amersham ImageQuant 800 (Cytiva 29399482). Optical measurements for the Precision Red and NanoBiT assays were measured on a Cytation 5 Multimode Plate Reader (Biotek).

### Synthesis of degraders 8-19

#### ***Scheme S1***

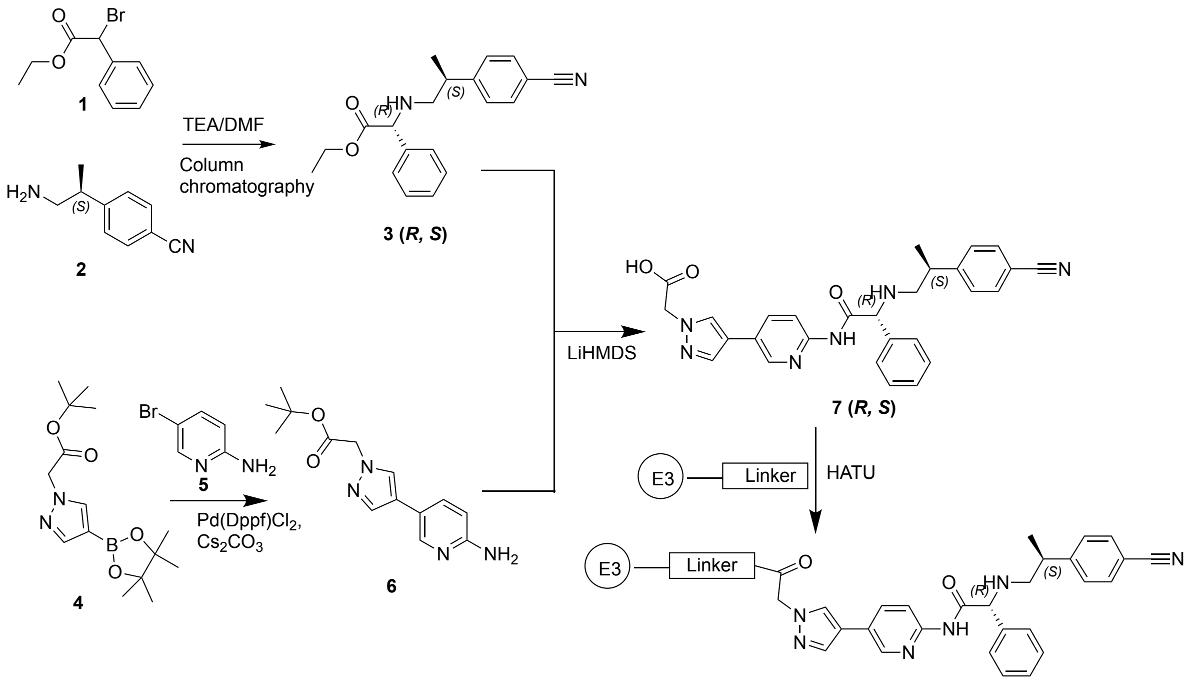


CPI-1612-based degraders (**8-14,19**)

#### ***Scheme S2***

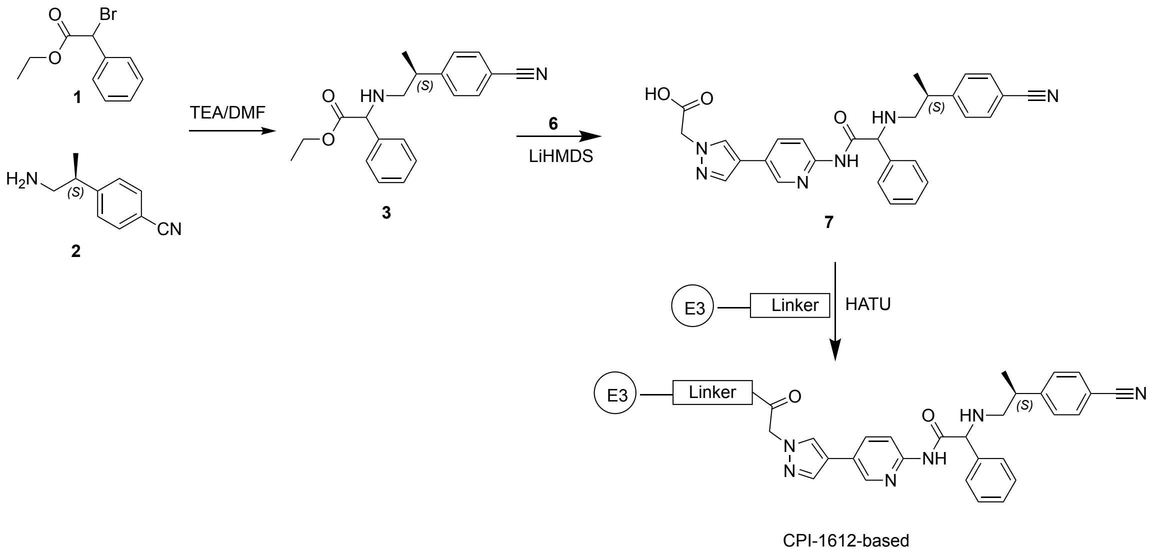


CPI-1612-based degraders (**15-18**)

### Synthetic procedures and compound characterization

Bifunctional compounds **8-19** were prepared according to Scheme S1 and Scheme S2.

***Synthesis of ethyl (R)-2-(((S)-2-(4-cyanophenyl)propyl)amino-2-phenylacetate; 3-(R,S) and 3-(S,S)***[2]

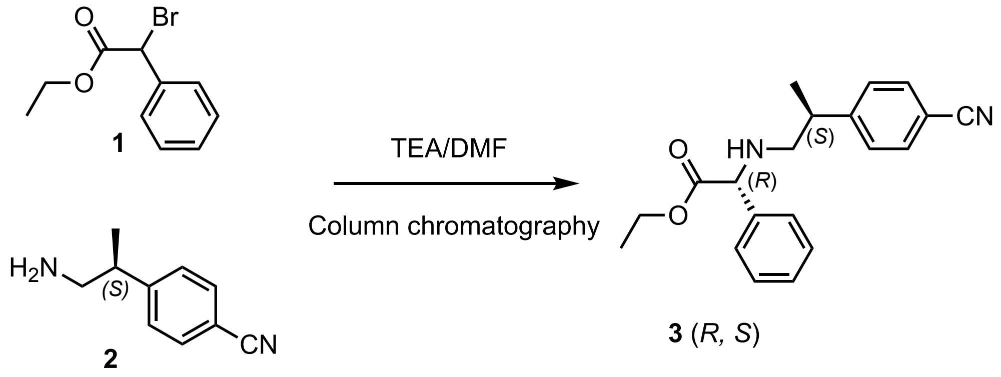


To the solution of (*S*)-4-(1-aminopropan-2-yl)benzonitrile hydrochloride (500 mg, 2.54 mmol) and ethyl 2-bromo-2-phenylacetate (773 mg, 560 µL, 3.18 mmol) in DMF, triethylamine (1.03 g, 10.2 mmol) was added. The reaction mixture was heated for 16 h at 90 °C with monitoring by TLC. After starting material was consumed, cold water (250 mL) was added, and the crude compound was extracted with ethyl acetate (2 x 250 mL). The organic fractions were pooled, dried on anhydrous $Na_2SO_4$, filtered, and then concentrated. The resulting residue was purified by silica gel chromatography (eluent: EA/Hexane, 4:6) to afford title compounds **3-(*S,S*) (**faster-eluting diastereomer, 300 mg/0.93 mmol (37%) and **3-(*R,S*)** (slower-eluting diastereomer, 368 mg/1.14 mmol, (45%)) as viscous oils. Scheme S1 proceeded from the pure compound while Scheme S2 proceeded from the unseparated mixture of diastereomers. **3-(*R,S*)**: $^1$H NMR (400 MHz, $CDCl_3$) δ 7.52 – 7.48 (m, 2H), 7.27 – 7.14 (m, 7H), 4.22 (s, 1H), 4.14 – 3.96 (m, 2H), 2.92 (h, *J* = 7.0 Hz, 1H), 2.69 (dd, *J* = 11.6, 6.2 Hz, 1H), 2.56 (dd, *J* = 11.6, 7.9 Hz, 1H), 1.18 (d, *J* = 6.9 Hz, 3H), 1.10 (t, *J* = 7.1 Hz, 3H). $^{13}$C NMR (101 MHz, $CDCl_3$) δ 172.66, 150.96, 132.49 (3C), 128.78(2C), 128.22(3C), 127.46(2C), 119.13, 110.37, 65.58, 61.36, 54.00, 40.62, 19.67, 14.18; **3-(*S,S*)**; $^1$H NMR (400 MHz, $CDCl_3$) δ 7.55 – 7.50 (m, 2H), 7.28 – 7.18 (m, 7H), 4.22 (s, 1H), 4.14 – 3.98 (m, 3H), 2.93 (h, *J* = 7.0 Hz, 1H), 2.75 (dd, *J* = 11.5, 7.9 Hz, 1H), 2.61 (dd, *J* = 11.5, 6.3 Hz, 1H), 1.19 (d, *J* = 6.9 Hz, 3H), 1.11 (t, *J* = 7.1 Hz, 3H). $^{13}$C NMR (101 MHz, $CDCl_3$) δ 172.85, 150.89, 132.50 (3C), 128.79(2C), 128.26(3C), 127.51(2C), 119.14, 110.42, 65.64, 61.34, 54.29, 40.70, 19.92, 14.20.

***Synthesis of tert-butyl 2-(4-(6-aminopyridin-3-yl)-1H-pyrazol-1-yl)acetate (6)***

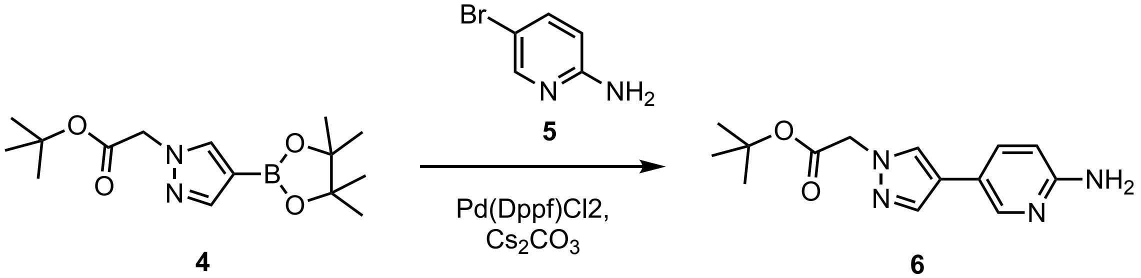


To the solution of 5-bromopyridin-2-amine **5** (2.00 g, 11.56 mmol) and tert-butyl 2-(4-(4,4,5,5-tetramethyl-1,3,2-dioxaborolan-2-yl)-1H-pyrazol-1-yl)acetate **4** (3.92 g, 12.7 mmol) cesium carbonate (11.3 g, 34.68 mmol)in dioxane/water (4:1), Pd(dppf)Cl2 (845.9 mg, 1.156 mmol) was added under the inert gas flow. The reaction mixture was heated for 16 h at 90 °C with monitoring by TLC. After completion, cold water (250 mL) was added, and the crude compound was extracted with ethyl acetate (2 x 250 mL). The organic fractions were pooled, dried on anhydrous $Na_2SO_4$, filtered, and then evaporated to get the crude compound. The product **6** was purified by column chromatography on silica gel (eluent: MeOH/DCM, 1:9) to obtain title compound as a brown liquid. Yield 2.23 g/8.13 mmol (70%); $^1$H NMR (400 MHz, cdcl$_3$) δ 8.18 (s, 1H), 7.70 (s, 1H), 7.61 (s, 1H), 7.56 – 7.46 (m, 1H), 6.52 (d, *J* = 8.4 Hz, 1H), 4.81 (s, 1H), 4.58 – 4.13 (m, 2H), 1.47 (s, 9H).

***Synthesis of 2-(4-(6-((R)-2-(((S)-2-(4-cyanophenyl)propyl)amino)-2-phenylacetamido)-pyridin-3-yl)-1H-pyrazol-1-yl)acetic acid. acetate (7)***

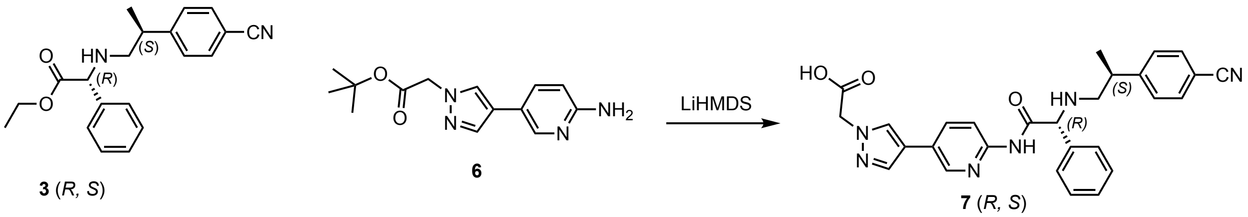


To tert-butyl 2-(4-(6-aminopyridin-3-yl)-1H-pyrazol-1-yl)acetate (340 mg, 1.24 mmol), and ethyl (*R*)-2-(((*S*)-2-(4-cyanophenyl)propyl)amino)-2-phenylacetate (**3-(*R,S*)**) (200 mg, 0.620 mmol) in anhydrous toluene (10 mL), lithium bis(trimethylsilyl)amide (311 mg, 1.86 mL, 1.0 molar, 1.86 mmol) was added at -78 °C. The reaction mixture was warmed to room temperature and stirred for 3 h with the reaction progress monitored by LCMS. After completion of the reaction, cold water (5 mL) was added, and the mixture was evaporated to afford the crude product as a brown solid. The product was purified by reverse phase column chromatography on C18 column (eluent: acetonitrile/water (0.05%TFA)) to obtain pure **7-(*R,S*)** as a pale yellow solid. 195 mg/0.39 mmol (64%); $^1$H NMR (400 MHz, DMSO) δ 11.41 – 11.26 (m, 1H), 9.76 – 9.42 (m, 1H), 8.60 (dt, *J* = 7.2, 1.6 Hz, 1H), 8.21 (d, *J* = 5.5 Hz, 1H), 8.06 – 8.00 (m, 2H), 7.96 (d, *J* = 5.2 Hz, 1H), 7.87 – 7.83 (m, 1H), 7.82 – 7.78 (m, 2H), 7.66 – 7.60 (m, 2H), 7.57 – 7.45 (m, 4H), 5.13 – 5.05 (m, 1H), 4.98 (d, *J* = 2.8 Hz, 2H), 3.28 (dd, *J* = 8.3, 5.7 Hz, 1H), 3.24 – 3.12 (m, 1H), 3.11 – 3.02 (m, 1H), 2.95

(d, $J$ = 8.9 Hz, 1H), 1.32 – 1.19 (m, 3H); HRMS (ESI): calcd for ($C_{28}H_{26}N_6O_3$ + $H^+$), 495.2145; found, 495.2141 $[M+H]^+$.

***General Amidation Method: A***

To the corresponding carboxylic acids (1.0 eq) dissolved in DMF (2-3 mL), HATU (1.25 eq) was added at RT. Subsequently, the corresponding alkyl amines (1.0 eq) and DIPEA (1.25 eq) were added, and the mixture stirred at RT for 14 h. The reaction progress was monitored by LCMS. After completion of the reaction, the products were purified by reverse phase chromatography on a Teledyne ISCO Combiflash instrument (C18 column, $H_2O$ and $CH_3CN$, 0.05% trifluoroacetic acid v/v, gradient: $H_2O$:$CH_3CN$ (90:10) to (0:100) over 18 min, elution time 7-8 min). Pure fractions were collected and combined and lyophilized for 16 h to afford bifuncionals **8**-**19**.

***Synthesis of (4R)-1-((S)-2-(8-(2-(4-(6-((R)-2-(((S)-2-(4-cyanophenyl)propyl)amino)-2-phenyl-acetamido)pyridin-3-yl)-1H-pyrazol-1-yl)acetamido)octanamido)-3,3-dimethylbutanoyl)-4-hydroxy-N-((S)-1-(4-(4-methylthiazol-5-yl)phenyl)ethyl)pyrrolidine-2-carboxamide (MC1, 10)*** The synthesis of MC-1 (**10**) employed general amidation method (A) to yield a colorless solid, yield 56 mg/0.53 mmol (65%); $^1$H NMR (400 MHz, DMSO) δ 10.41 (s, 1H, amide rotamer), 10.25 (s, 1H, amide rotamer), 8.91 (s, 1H), 8.50 (ddd, $J$ = 5.4, 2.3, 0.9 Hz, 1H), 8.29 (d, $J$ = 7.8 Hz, 1H), 8.11 (d, $J$ = 1.8 Hz, 1H), 8.04 (t, $J$ = 5.6 Hz, 1H), 7.95 (dd, $J$ = 8.7, 5.8 Hz, 1H), 7.92 – 7.83 (m, 2H), 7.75 – 7.65 (m, 3H), 7.42 – 7.14 (m, 12H), 5.02 (d, $J$ = 3.5 Hz, 1H), 4.91 – 4.79 (m, 1H), 4.71 (s, 2H), 4.49 – 4.32 (m, 3H), 4.21 (s, 1H), 3.54 (d, $J$ = 4.0 Hz, 2H), 3.05 – 2.91 (m, 3H), 2.60 (dd, $J$ = 14.3, 7.2 Hz, 3H), 2.38 (s, 3H), 2.17 (dt, $J$ = 14.8, 7.6 Hz, 1H), 2.04 (dt, $J$ = 14.1, 7.1 Hz, 1H), 1.94 (t, $J$ = 10.9 Hz, 1H), 1.72 (ddd, $J$ = 12.9, 8.5, 4.7 Hz, 1H), 1.50 – 1.37 (m, 3H), 1.30 (d, $J$ = 7.0 Hz, 3H), 1.23 – 1.09 (m, 9H), 0.86 (s, 9H). $^{13}$C NMR (101 MHz, dmso) δ 172.48, 171.85, 171.06, 170.06, 166.62, 152.16, 151.91, 149.82, 148.20, 145.11, 144.77, 139.79, 136.85, 134.83, 132.71, 132.65, 131.56, 130.13, 129.26 (3C), 129.02, 128.76 (2C), 127.96, 127.72 (2C), 126.82 (3C), 125.05, 119.49, 118.81, 113.69, 109.35, 69.20, 66.24, 58.98, 56.78, 54.67, 54.37, 48.13, 39.15, 38.18, 35.63, 35.32, 29.37, 28.89, 26.89 (3C), 26.72, 25.81, 22.87, 19.73, 16.43. HRMS (ESI): calcd for ($C_{59}H_{71}N_{11}O_6S$ + $H^+$), 1062.5388; found, 1062.5371 $[M+H]^+$.

**Primary degrader screen**

HAP-1 cells were plated at 3 x $10^5$ cells/well in 2 mL IMDM in 6-well plates. After ~24 h, media was aspirated and 1700 µL fresh media was replaced. Cells were treated dropwise with 300 µL of **8-19**, dCBP-1, or JQAD-1-containing media (0.1-0.2% final concentrations DMSO). Control wells were treated with DMSO at the same 0.1 or 0.2% final concentration of a given compound. Following an approximately 24 h incubation, media was aspirated, cells were washed once with 1 mL ice-cold PBS, and lysed in 150 µL 1x RIPA buffer containing 1x protease inhibitor cocktail. Lysates were incubated on ice for 15-30 min before clarification by centrifugation at 14,000 rpm for 15 min at 4°C. Protein quantification was performed with the Precision Red Assay according to manufacturer's protocols. 15 µg of total lysate were loaded per well on gels for **8-19** and JQAD-1, while 14 µg of total lysate were loaded per well for dCBP-1 due to samples with slightly lower protein concentrations.

**Quantification of $D_{max}$**

Western blot densitometry analysis was performed using ImageJ software to quantify relative maximum degradation ($D_{max}$) values from the primary degrader screen Western blots and Fig. S1 (n = 1). Area under the curve (AUC) from images in RGB format were determined using the gel band densitometry functions on ImageJ for the indicated antibody and exposure time, for EP300, CREBBP, and Vinculin. Band intensities of EP300 or CREBBP were normalized to Vinculin intensities on the same blot as well as to the DMSO control. Subtraction of relative intensities from the DMSO control yielded relative degradation percentages, and the highest degradation percentage is reported as $D_{max}$.

**Conjugation of CPI-1612-C6-PEG5-biotin to streptavidin magnetic beads**

CPI-1612 coated bead was prepared using Thermo Scientific™ Pierce™ Streptavidin Magnetic Beads (Thermo Scientific, 88817). Briefly, 20 mM CPI1612-C6-biotin was prepared in DMSO. The streptavidin magnetic beads were prewashed with 0.1% NP-40 1×PBS (v/v) 3 times using a magnetic racker. Then, 20 mM CPI1612-C6-biotin, 0.1% NP-40 1×PBS and 10 mg/mL prewashed beads were mixed at a ratio of 1:40:22 (v/v/v), and incubated at 4 °C overnight while rotating. The next day, this coating system was incubated at RT for 1 h by rotating before wash. Beads were washed with a series buffer: 1 mL high salt buffer (0.1% NP-40, 200 mM NaCl, 1×PBS) 2 times, 1 mL $H_2O$ 2 times, and 1 mL 0.1% NP-40 1×PBS 2 times. Supernatants were removed and 0.1% NP40 1×PBS added to the beads for a final concentration of 10 mg/mL.

**Competitive affinity capture and immunoaffinity profiling**

Hela nuclear extracts (IPRACELL CC-01-20-50) were first diluted to a working protein concentration of 1 mg/mL with high salt buffer (0.1% NP-40, 200 mM NaCl, 1×PBS) containing 1× protease inhibitors (Cell Signaling Technology 5871) and 10 µM MG132 (Selleckchem,

S2619). The lysates were centrifuged at 20 000 g, 4°C for 30 mins, and then aliquoted 350 μL supernatant into individual tubes, each tube corresponding to different conditions (i.e. lanes on gel). 0.35 μL 0.5, 1, 5 mM CPI-1612 (1000x) or MC-1 (1000x) was added to 350 μL proteome (working concentration of competitor: 0.5, 1, 5 μM), vehicle group added 0.35 μL DMSO instead. Samples were pre-incubated at 4 °C for 1.5 hrs while rotating. After pre-incubation, 52.5 μL coated beads were added to competitor/vehicle treated lysates. These mixtures were rotated overnight at 4°C. The magnetic capture beads were then subjected to a series of washes (1 mL×2 times) using high salt buffer (0.1% NP-40, 200 mM NaCl, 1×PBS) followed by 1×PBS. After the last wash, discarded all the supernatants and added 50 μL 1 x loading buffer (Invitrogen, NP0007) containing 100 mM DTT. Resuspended the beads, boil them at 95 °C for 10 mins. Transfer the supernatant to a new tube after beads absorbed by the magnetic racker. Did the elution again in the same way and combine the eluted samples. For immunoblot analysis, 15 μL of each eluted sample was loaded onto a 3-8% Tris-Acetate gel (Invitrogen, EA03755BOX) to proteome separation and transferred to western blot at 30 V for 1 hour by using 0.45 μM nitrocellulose membrane (Invitrogen, LC2001). Membranes were blocked, probed with anti-P300 antibodies (Santa Cruz Biotech, SC-584, 1:1000 dilution), anti-CREBBP antibodies (Cell Signaling, 7389S, 1:1000 dilution), and anti-vinculin antibodies (Cell Signaling, 18799S, 1:1000 dilution), then washed and developed according to the manufacturers protocol. Quantification of affinity capture of EP300/CREBBP was did by ImageJ and data was processed in GraphPad Prism 10 by using non-linear regression model.

**Mechanistic Analyses**

*Dose-dependency in HAP1 cells*

HAP-1 cells were plated at 4 x $10^5$ cells/well in 2 mL media. After 16-24 h, cells were treated with MC-1 as described in the “Primary Degrader Screen” section. All wells had 0.1% final concentrations DMSO. Following a 6 h incubation, cells were washed, lysed in RIPA, and quantified as described above. 15 μg of total lysate were loaded per well on gels.

*Time-dependency and co-incubation with VH298*

HAP-1 cells were plated at 4 x $10^5$ cells/well in 2 mL media. After 24 h, cells were treated according to the method described above with a 1 μM final concentration of MC-1, and for the VH298 co-incubation samples only, simultaneously treated with a 10 μM final concentration of VH298 (Tocris 6156). Final DMSO concentrations were 0.1% for the time-course samples and 0.2% for the VH298 co-incubation. Cells were lysed at 0 (no treatment), 1, 3, 6, and 24 h following treatment. The time-dependency and co-incubation experiments were performed at the same time and samples run on the same blots.15 μg of total lysate were loaded per well for the CREBBP blot, and 25 μg of total lysate were loaded per well for the EP300 blot.

*Pre-incubation with CPI-1612*

HAP-1 cells were plated at 4 x $10^5$ cells/well in 2 mL media. After 24 h, cells were treated as described above with a final concentration of 1µM CPI-1612 (0.1% final DMSO concentration) or DMSO only (0.1% final concentration) and pre-incubated for 1 h. After 1 hour, 2 µL of a 1mM MC-1 stock in DMSO were directly added to the wells (final DMSO concentrations = 0.2%). Cells were lysed at 1, 3, 6, and 24 h following MC-1 addition, with the MC-1 only control lysed at 24 h after MC-1 addition. 15 µg of total lysate were loaded per well on gels.

*6h treatment of All VHL and CRBN-binding compounds in HAP-1 cells*

HAP-1 cells were plated at 5 x $10^5$ cells in 2 mL media per well in a 6-well dish. After 16-24 h, cells were treated according to the method described above with the indicated final concentrations of **8-19** (final DMSO concentrations = 0.1%), or DMSO vehicle (0.1% final concentration). Cells were lysed after 6 h as described above. 12 µg of total lysate were loaded per well on gels for the blot with VHL-binding compounds and 15 µg of total lysate per well for the blot with CRBN-binding compounds.

*JQAD-1 6h treatment in HAP-1 cells*

HAP-1 cells were plated at 5 x $10^5$ cells in 2 mL media per well in a 6-well dish. After 16-24 h, cells were treated according to the method described above with the indicated final concentrations of JQAD-1 (final DMSO concentrations = 0.1%), or DMSO vehicle (0.1% final concentration). Cells were lysed after 6 h as described above. 15 µg of total lysate were loaded per well on gels.

**Histone modification analyses**

HAP-1 cells were plated at 5 x $10^5$ cells in 2 mL media per well in 6-well dishes. MCF-7 cells were plated at 4 x $10^5$ cells/well in 2 mL media per well in 6-well dishes. After 16-24 h cells were treated as described above with MC-1 or CPI-1612-containing media (0.1% final concentrations of DMSO). All cells were treated in duplicate for 6 h, with one set harvested in RIPA as described above for total protein blots and the other harvested in parallel as described below for histone analyses.

For histone modification analyses, after 6 h, cells were washed with 1 mL ice-cold PBS and either scraped in 1 mL ice-cold PBS, pelleted (500 rcf, 4 °C, 5 min), media aspirated, and flash frozen and stored at -80 °C before later resuspension and lysis as described below. Alternatively, cells were washed once with 1 mL ice-cold PBS and lysed directly in the plate in 150 µL of Nuclear Isolation Buffer (NIB)[3] (15 mM pH 7.5 Tris-HCl, 60 mM KCl, 15 mM NaCl, 5 mM $MgCl_2$, 1 mM $CaCl_2$, 250 mM sucrose, 1X protease inhibitor cocktail [Cell Signaling Technology #5871]), 1 mM DTT, and 10 mM sodium butyrate) with 0.1% IGEPAL. Suspensions of lysed cells were transferred into tubes on ice and incubated for $\geq$5 min. Nuclei were pelleted (600 rcf, 4 °C, 5 min) and the supernatant discarded. Pelleted nuclei were subjected to two cycles of a wash with 150 µL NIB (without IGEPAL CA-630) followed by centrifugation (600 rcf, 4 °C, 5 min) to remove all IGEPAL CA-630. Nuclei were re-suspended in 400 µL 0.4 N $H_2SO_4$ and rotated at 4 °C for 16 h. The following day, samples were centrifuged (11,000 rcf, 4 °C, 10 min), supernatants transferred into new tubes (pellets discarded). 100 µL of 100% TCA was added per sample (final TCA concentration: 20%) and tubes were inverted once. Histones were precipitated at 4°C for 16-24 h or for ≥ 4 h on ice, then centrifuged (11,000 rcf, 4 °C, 5 min) and supernatants discarded, with histones now visible as films on the sides and at the bottom of tubes. Histones were subjected to two cycles of washing (1 mL of ice-cold acetone + 1% 1M HCl, followed by 1 mL of 100% ice-cold acetone) followed by centrifugation (11,000 rcf, 4 °C, 5 min). Samples were air dried at room temperature and resuspended in 50 µL of $ddH_2O$. Protein was quantified with the Precision Red assay and 1 or 2 µg were loaded on per well of each gel (exact amounts indicated in captions of uncropped Western blots).

**Quantitative proteomics analysis of MC-1 targets**

HAP-1 cells were plated at 4 x $10^6$ cells in 10 mL media in 10-cm dishes. After 24 h, media was aspirated and replaced with 10 mL media containing 1 µM MC-1, 1 µM dCBP-1, or DMSO vehicle (all DMSO final concentrations = 0.01%). After 6 h of treatment, media was aspirated, cells were washed in 5 mL cold PBS, scraped in 5 mL cold PBS, pelleted (500 rcf, 4 °C, 5 min), re-suspended in 1 mL PBS, transferred to 1.7 mL tubes, and pelleted (500 rcf, 4 °C, 5 min), followed by a dry spin (500 rcf, 4 °C, 5 min). Cell pellets were flash frozen and stored at -80 °C.

Frozen pellets of HAP-1 cells treated with DMSO, MC-1, and dCBP-1 were suspended in 400 µL of EasyPep lysis buffer (Thermo Scientific, A45735) containing 10U of Pierce® Universal Nuclease (Thermo Scientific, 88700) and sonicated using a probe sonicator (QSonica). The protein concentrations were determined using Pierce® BCA assay kit (Thermo Scientific, 23225) and 20 µg of each sample was digested in 200 µL of 1:1:1:1 100 mM HEPES/lysis buffer/reducing solution/alkylating solution from the EasyPep™ MS Sample Prep Kit (Thermo, A40006) containing 1 µg Trypsin/LysC protease mix (Thermo, A40007) overnight with shaking at 37 °C. 15-plex reagents (125 µg, Thermo, A52045) were dissolved in 20 µL acetonitrile (ACN) and added into each sample, followed by incubation at RT for 1 hr and quenched with 50 µL 20% formic acid + 5% hydroxylamine for 15 mins at RT. The labeled peptides were pooled and cleaned up using EasyPep Mini columns (Thermo Scientific, A40006) by following the provided protocol. The peptides were eluted from the column and dried in a speed-vac.

The dried peptide sample was dissolved in 0.1% formic acid and subjected to an offline high pH reversed-phase fractionation using a Waters Acquity UPLC system coupled with a fluorescence detector (Waters) and an Xbridge Peptide BEM™ 2.5 µm C18 column (150 mm x 3.0 mm, Waters) operating at 0.35 mL/min. The peptides were separated using mobile phase solvent A (10 mM ammonium formate pH 9.4) and mobile phase solvent B (90% acetonitrile in 10 mM ammonium formate pH 9.4). with the following gradients: 6% solvent B (0-1 min), 6-10% solvent B (1-1.5 min), 10-50% solvent B (1.5-60 min), 50-90% solvent B (60-65 min), 90-6% solvent B (65-70 min). The fractions were collected every minute generating 60 fractions and pooled into 12 fractions by concatenation. The peptide fractions were dried and reconstituted in 0.1% formic acid (FA).

Each of the peptide samples was loaded onto a Dionex U3000 RSLC in front of an Orbitrap Eclipse (Thermo) equipped with an EasySpray ion source with mobile phases Solvent A consisted of 0.1% FA in water and Solvent B of 0.1% FA in 80% acetonitrile in water. The loading pump consisted of Solvent A operated at 7 µL/min for the first 6 min then reduced to 2 µL/min when the valve was switched to bring the trap column (Acclaim™ PepMap™ 100 C18 HPLC column, 3 µm, 75 µm i. d., 2 cm, part no. 164535). The gradient pump was set to a flow rate of 300 nL/min and used a linear LC gradient of 5-7% B for 1 min, 7-30% B for 134 min, 30-50% B for 35 min, 50-95% B for 4 min, holding at 95% B for 7 min, then re-equibiliration to 5% B for 17 min. All runs used the TopSpeed method with 3 sec cycle time that consisted of spray voltage at 1800 V and ion transfer temperature of 275 °C. MS1 scans were acquired in the Orbitrap with a 120,000 resolution, standard 100% AGC target, max injection time of 50 ms, and mass range set to 400-2000 *m/z*; MS2 scans were acquired in the Orbitrap as well using Turbo TMT method with a resolution of 30,000, standard 100% AGC target, max injection time of 54 ms, HCD energy of 38%, isolation window of 0.4 Da, minimum intensity set at 2.5e4, and charges 2-6 for MS2 selection. Advanced Peak Determination, Monoisotopic Precursor Selection (MIPS), and EASY-IC for internal calibration were all enabled with dynamic exclusion set to a count of 1 for 15 sec.

All .raw file from the 12 injections were pooled together as fractions for one experiment and searched with Proteome Discoverer 2.4 using Sequest. The data was searched against the nonredundant UniProt Human database (accessed in November 2023) using a full tryptic digest, 2 max missed cleavages, with 6 and 40 amino acids minimum and maximum peptide lengths, respectively. The MS1 precursor mass tolerance was set to 10 ppm, MS2 fragment tolerance at 0.02 Da, variable oxidation on methionine (+15.995 Da), and fixed modifications of carbamidomethyl on cysteine (+57.021 Da), TMTpro (+304.207 Da) lysine and peptide N-terminus. FDR analysis was done using Percolator and TMTpro reporter ions were quantified using the Reporter Ion Quantifier node using the unique peptides or unique + razor peptides with normalization using total peptide intensity across all channels. The TMTpro channel assignments can be found in the Supplementary Table S1.

**Cytotoxicity analyses (HAP-1 and MCF-7)**

MC-1, dCBP-1, and CPI-1612 were analyzed for growth inhibition against HAP-1 and MCF-7 cells using CellTiter-Glo Luminescent Cell Viability Assay (Promega, G7572). Briefly, 3 x $10^3$ MCF-7 cells or 2 x $10^3$ HAP-1 cells were plated per well in white-walled 96-well plates (Corning 3610) and allowed to adhere for 24 h. Then cells were treated with different concentrations of MC-1, dCBP-1or CPI-1612 (25 µM, 8.333 µM, 2.778 µM, 0.926 µM, 0.309 µM, 0.103 µM, 0.034 µM, 0.011 µM, with 0.25% (v/v) maximum final concentrations of DMSO) for 96 h. Vehicle control groups were treated with 0.25% (v/v) final concentrations of DMSO. Experiments were performed in quadruplicate. After 96 h incubation, cell viability was determined using the CellTiter-Glo

Luminescent Cell Viability Assay according to the manufacturer's instructions. Luminescence was measured on a BioTek Synergy 2 plate reader and resulting data was reported as normalized percent cell viability, with all values normalized to the DMSO vehicle controls. Results were plotted and half-maximal inhibition values were calculated from the nonlinear fit of dose-response data in GraphPad Prism 10.

### Full Blots for Fig. 2B (Primary Degrader Screen)

### **8** & **14**

EP300 (**14** = lanes 1-6, **8** = 7-12)

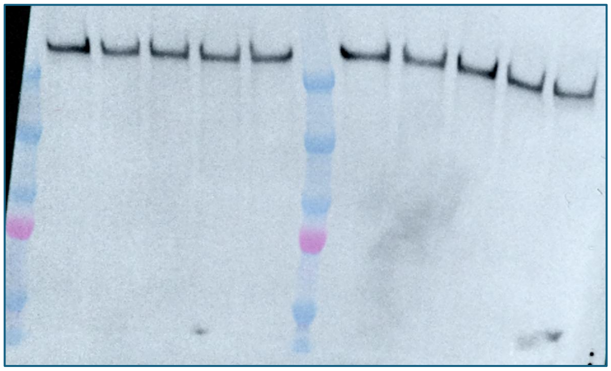

CREBBP (**14** = lanes 1-6, **8** = 7-12)

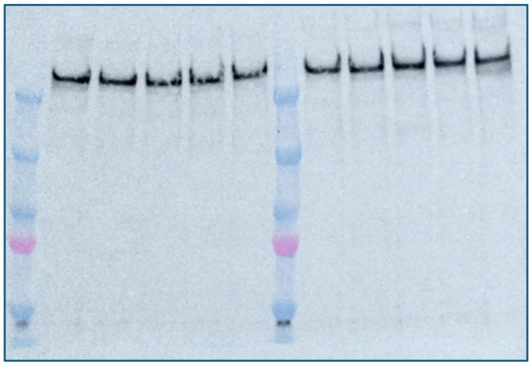

Vinculin (**14** = lanes 1-6, **8** = 7-12)

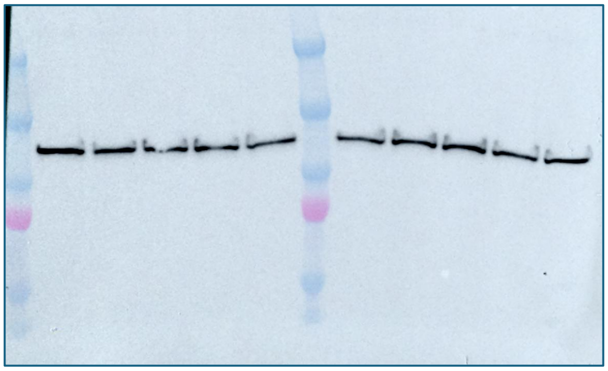

### **9**

EP300

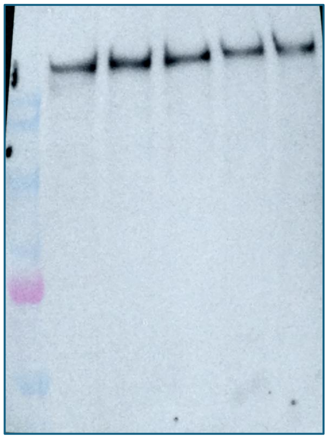

CREBBP

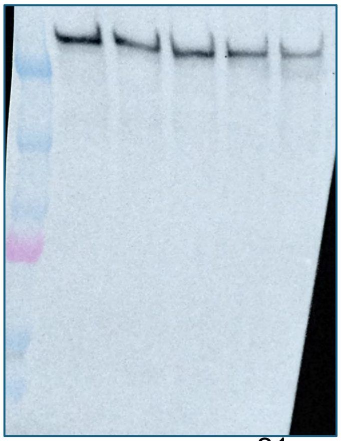

Vinculin

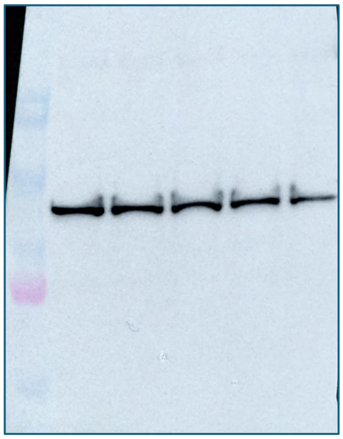

**10**

EP300 (lanes 1-6)

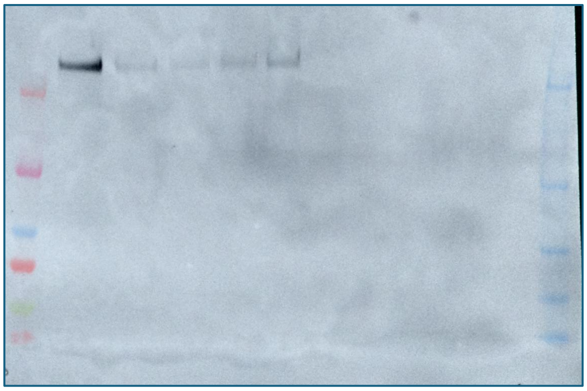

CREBBP (lanes 1-6)

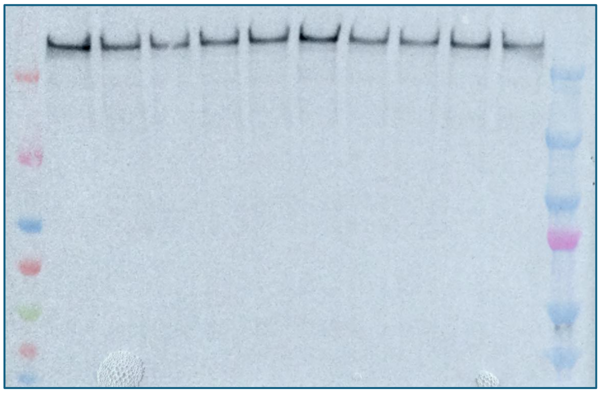

Vinculin (lanes 1-6)

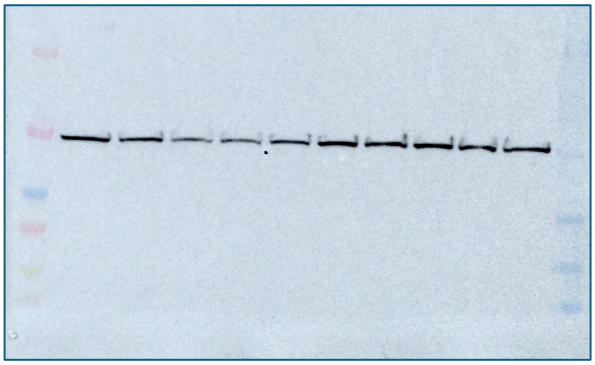

**11** & JQAD-1

EP300 (**11** = lanes 7-12, JQAD1 = 1-6)

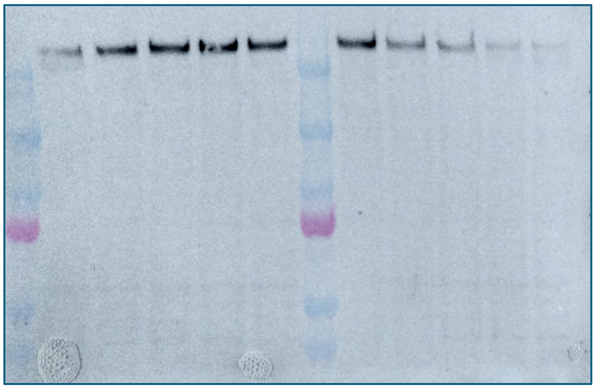

CREBBP (**11** = lanes 7-12, JQAD1 = 1-6)

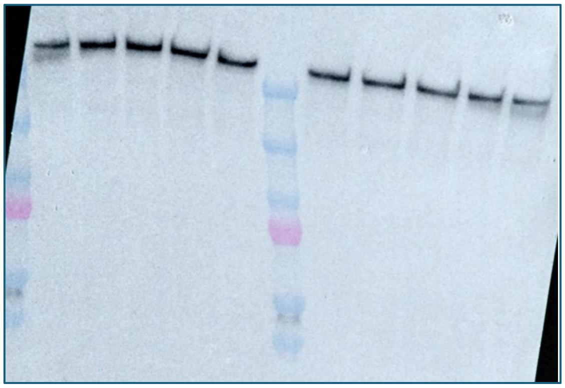

Vinculin (**11** = lanes 7-12, JQAD1 = 1-6)

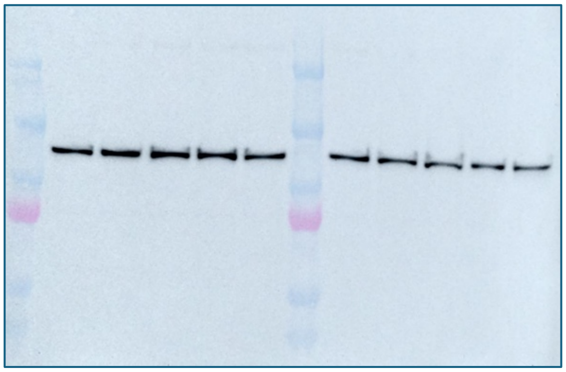

**12 & 13**

EP300 (**13** = lanes 1-6, **12** = 7-12)

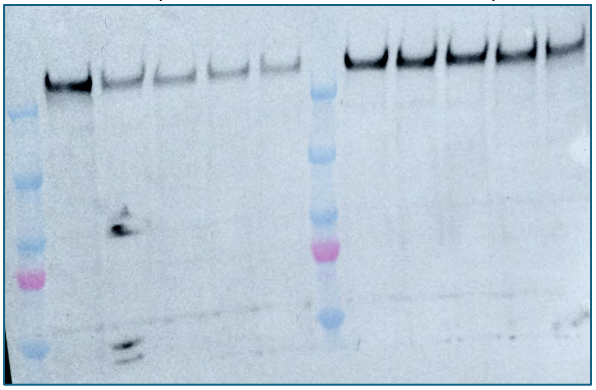

CREBBP (**13** = lanes 1-6, **12** = 7-12)

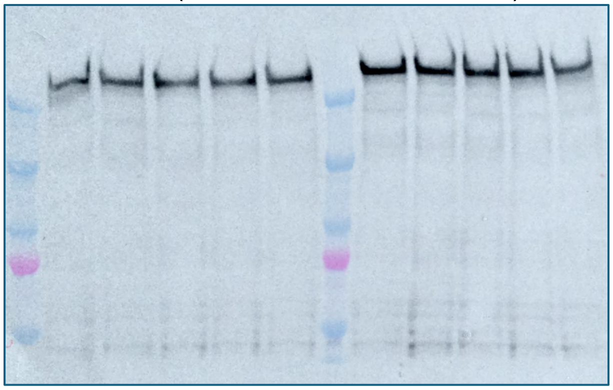

Vinculin (**13** = lanes 1-6, **12** = 7-12)

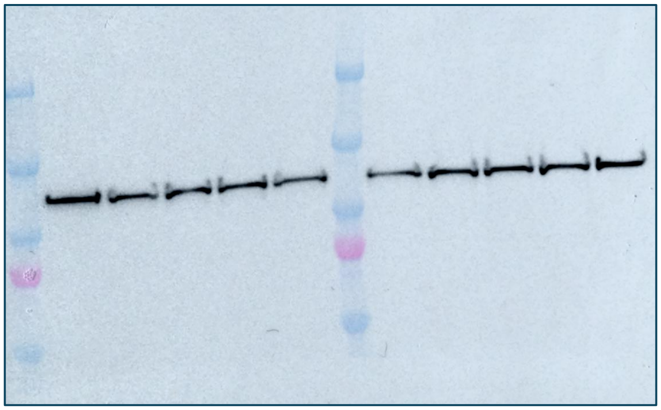

**15** & **16**

EP300 (**16** = lanes 1-6, **15** = 7-12)

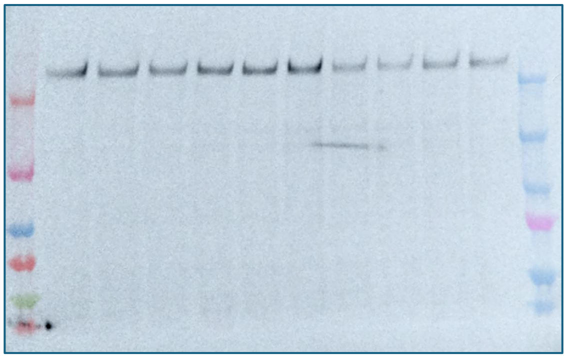

CREBBP (**16** = lanes 1-6, **15** = 7-12)

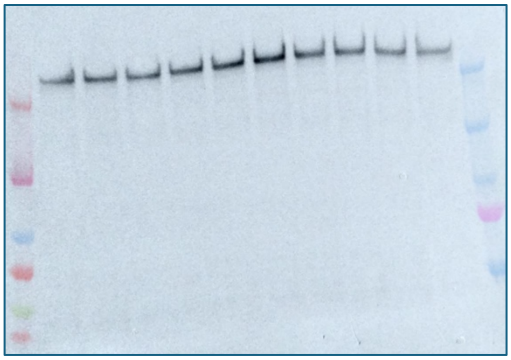

Vinculin (**16** = lanes 1-6, **15** = 7-12)

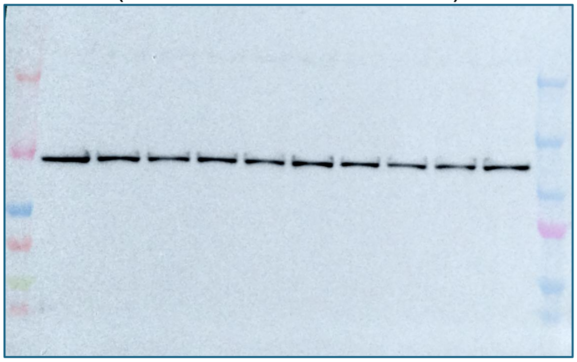

**17**

EP300

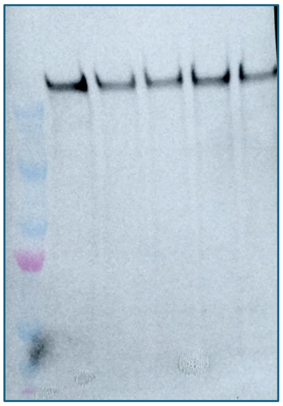

CREBBP

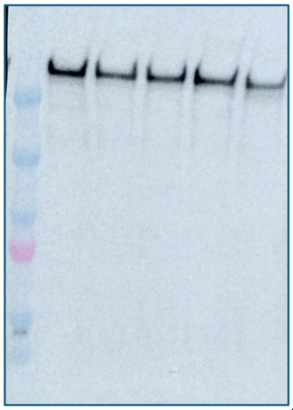

Vinculin

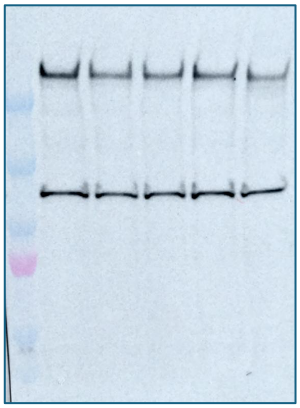

**18**

EP300

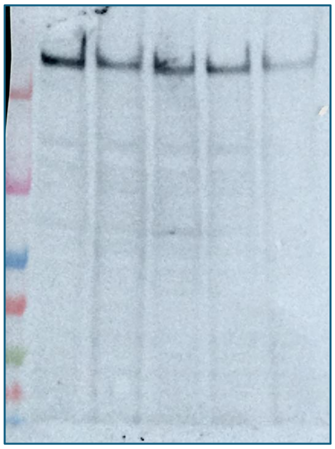

CREBBP

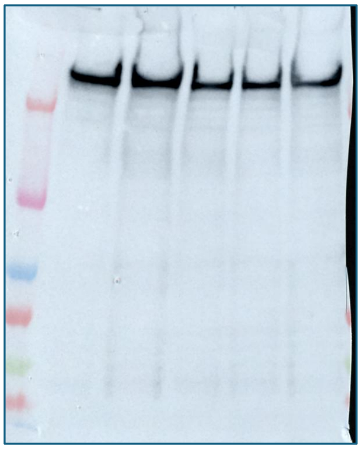

Vinculin

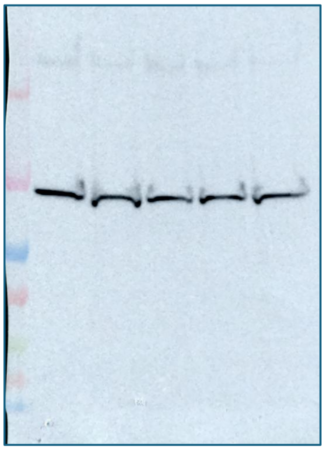

**19**

EP300 (lanes 1-6)

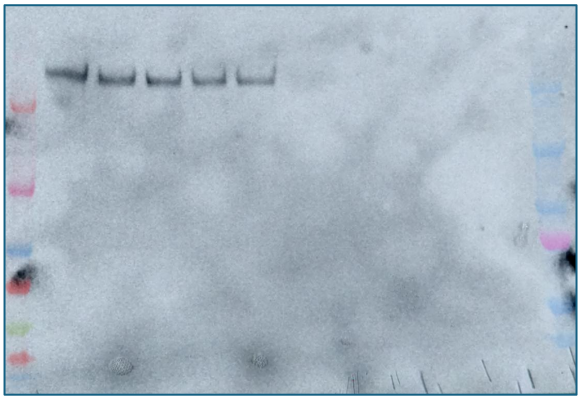

CREBBP (lanes 1-6)

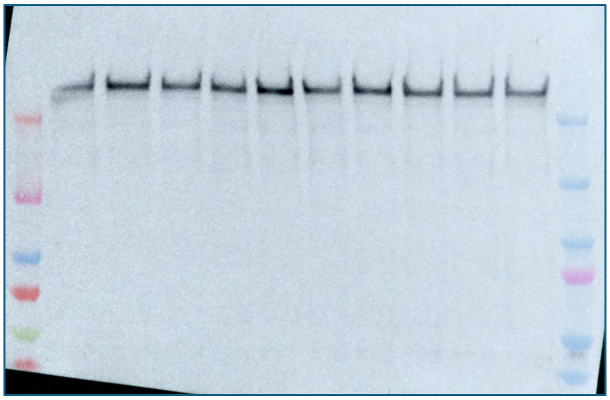

Vinculin (lanes 1-6)

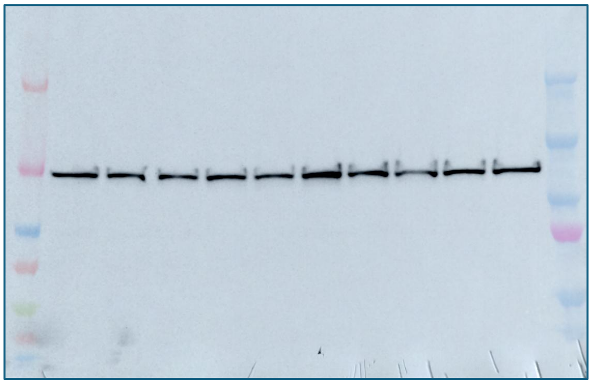

**Full Blots for Fig. 3D (MC-1 Dose-Dependence)**

EP300

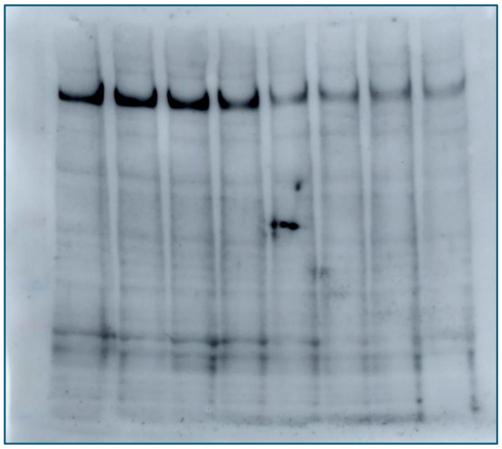

CREBBP

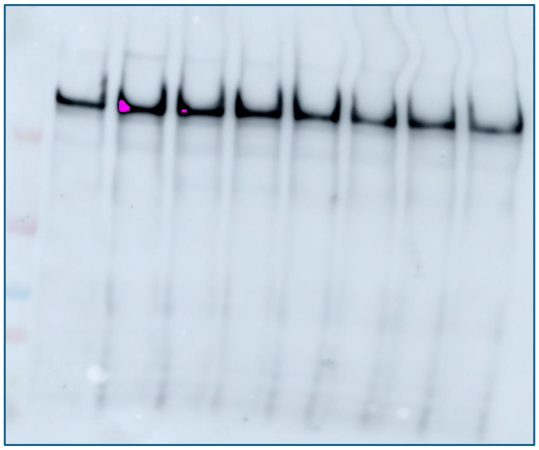

Vinculin

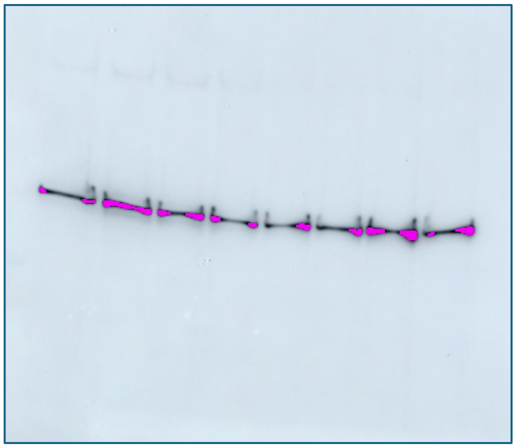

Vinculin

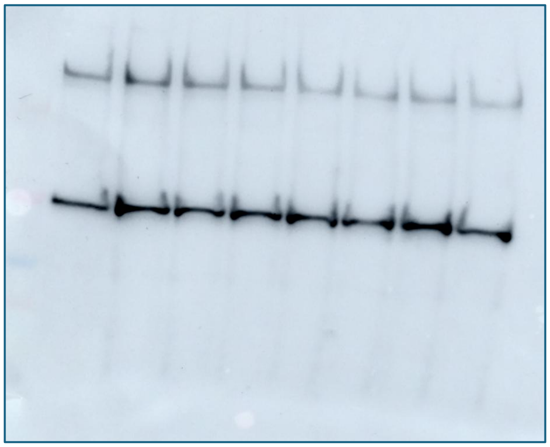

**Full Blots for Fig. 3 E & F (Time-course & VH298 Co-incubation)**

EP300

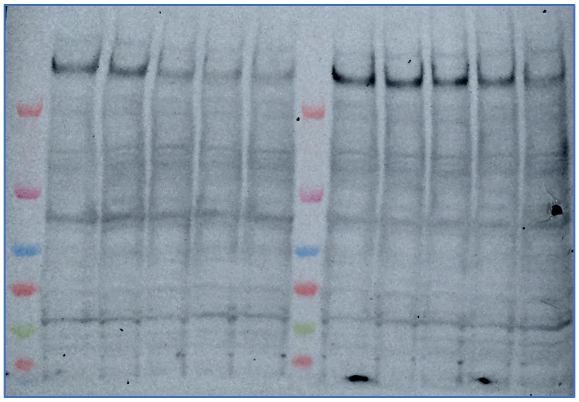

CREBBP

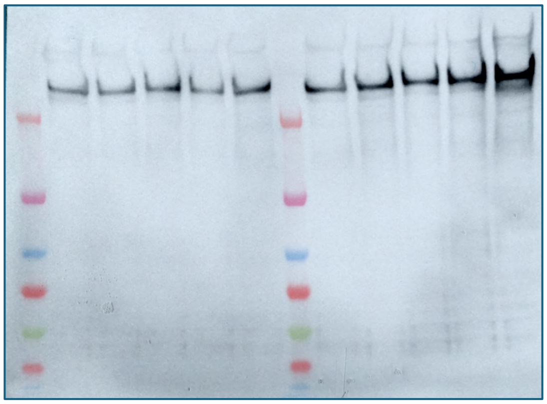

Vinculin

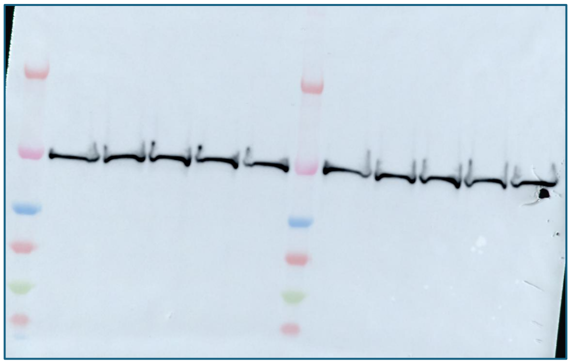

Vinculin

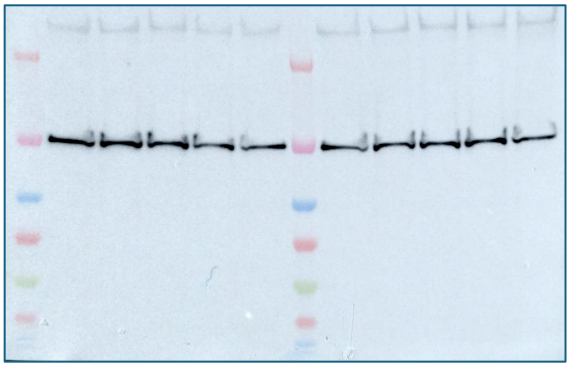

**Full Blots for Fig. 3G (CPI-1612 Pre-incubation)**

EP300 (lanes 3-8)

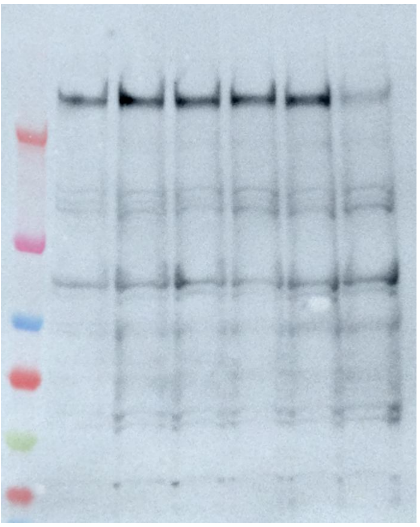

CREBBP (lanes 3-8)

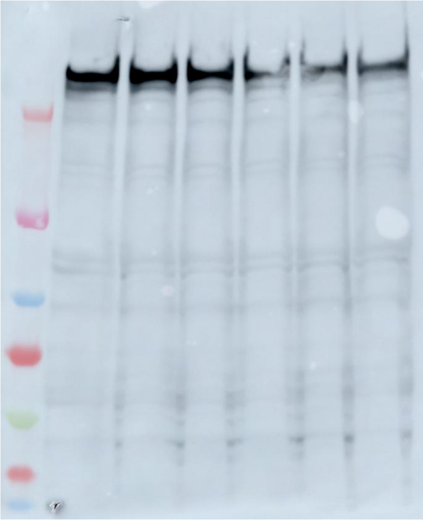

Vinculin (lanes 3-8)

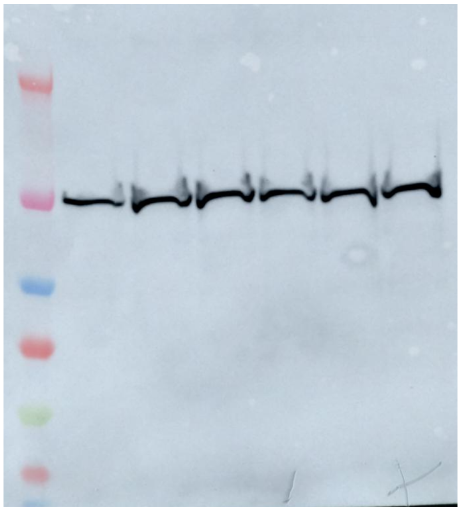

Vinculin (lanes 3-8)

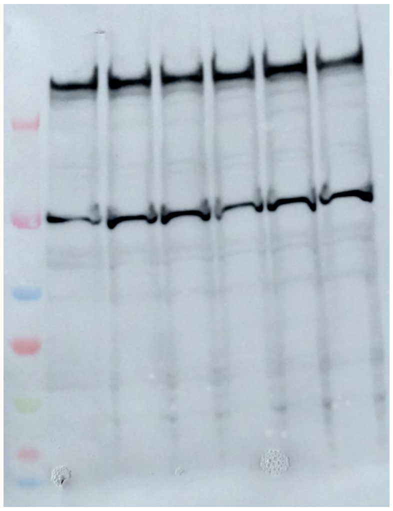

**Full Blots for Fig. 5A & S3E (Histone Acetylation in HAP1)**

H3K18Ac (lanes 1-6 = MC-1, 7-12 = CPI)

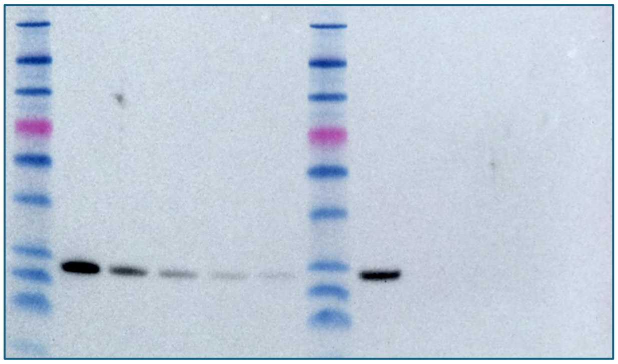

H3K23Ac

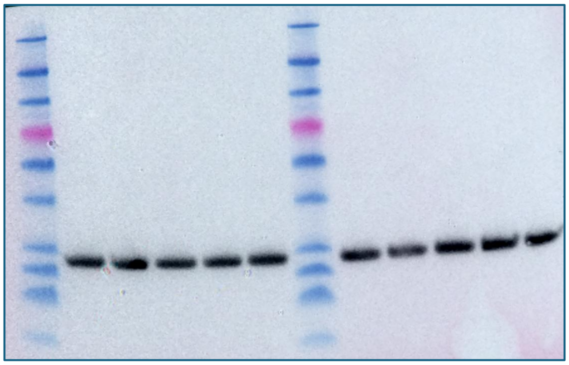

EP300

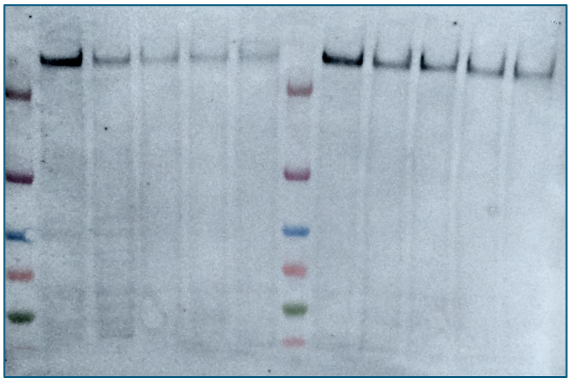

CREBBP

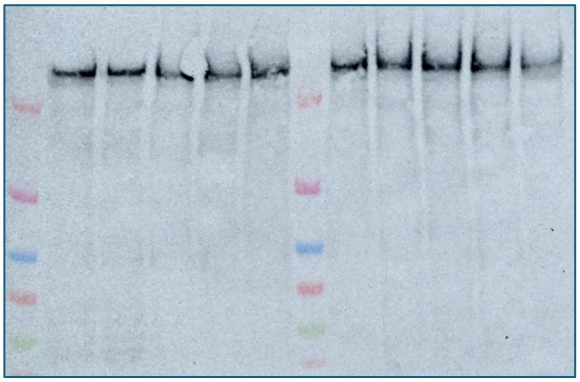

Vinculin

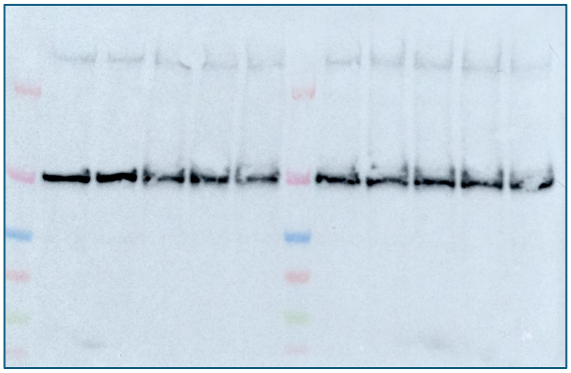

**Full Blots Fig. 5B (Histone Acetylation in MCF-7)**

EP300 (lanes 7-12)

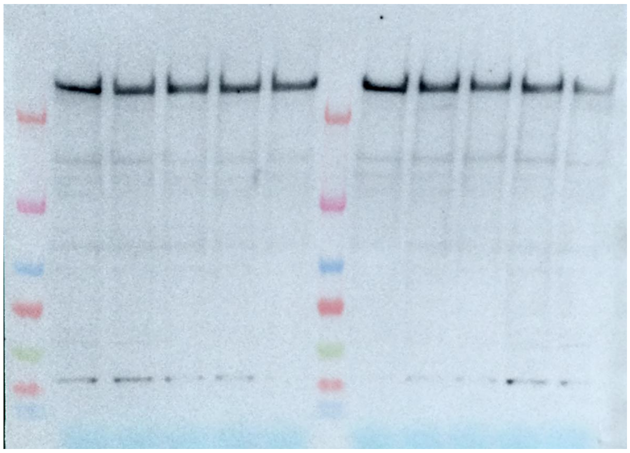

Vinculin (lanes 7-12)

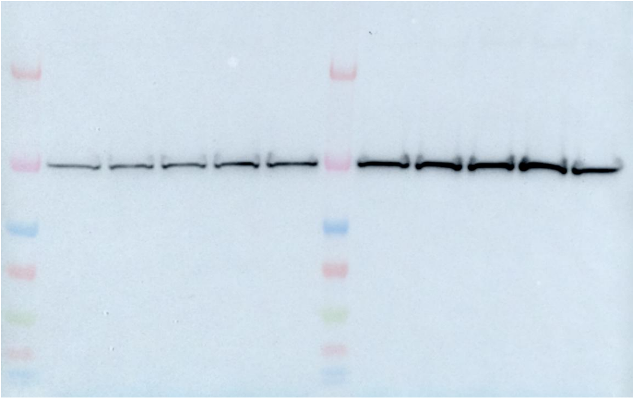

H3K18Ac (lanes 1-7)

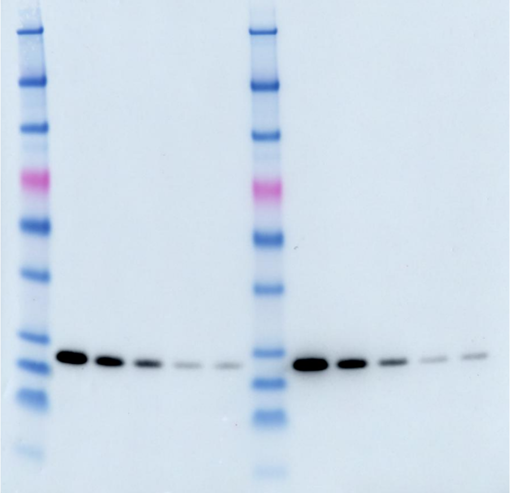

**Full Blots for Fig. S1 (JQAD-1 Dose-Dependence)**

EP300 (lanes 1-7)

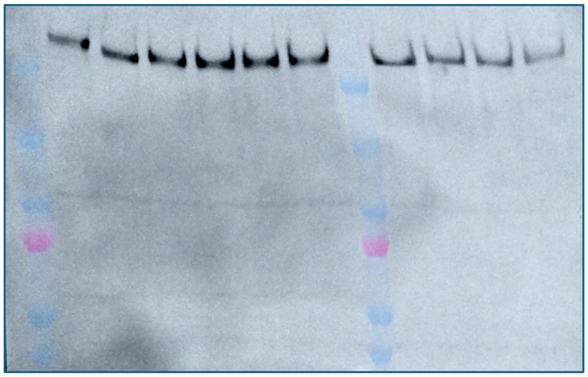

CREBBP

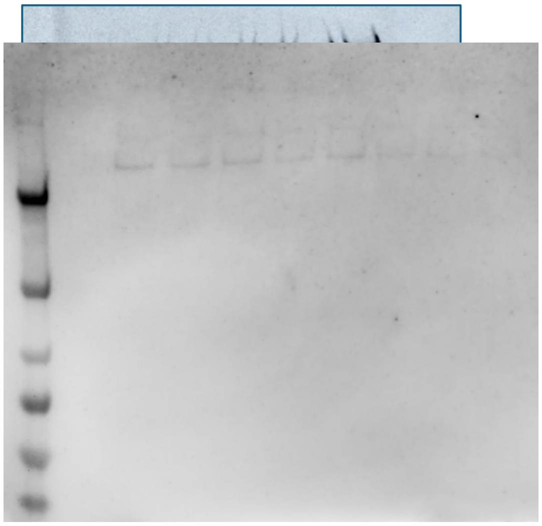

Vinculin (lanes 1-7)

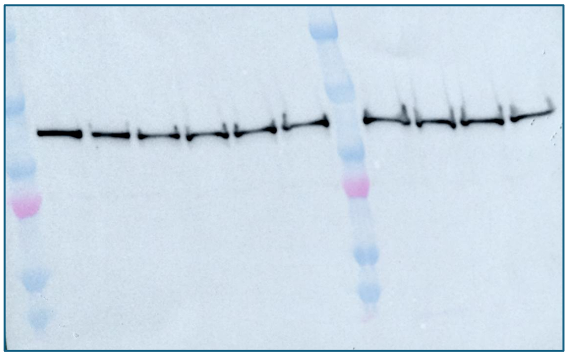

Vinculin

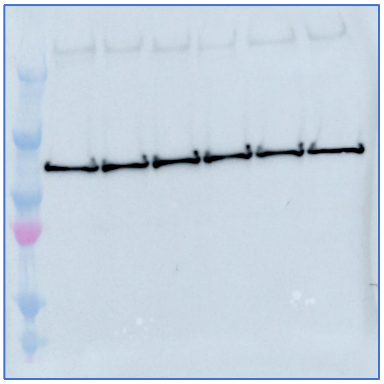

**Full Blots for Fig. S3C (CPI-1612 Affinity Capture)**

EP300 capture

EP300 input

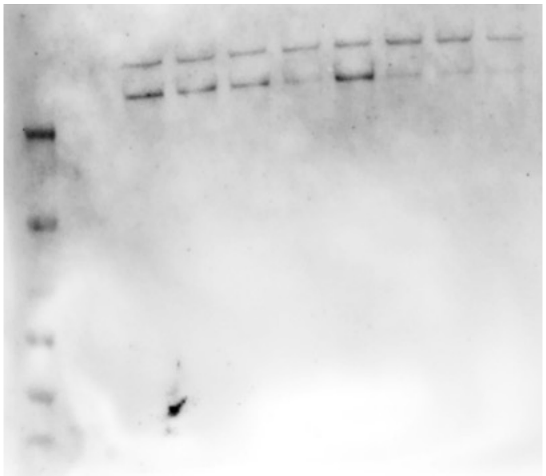

CREBBP capture

CREBBP input

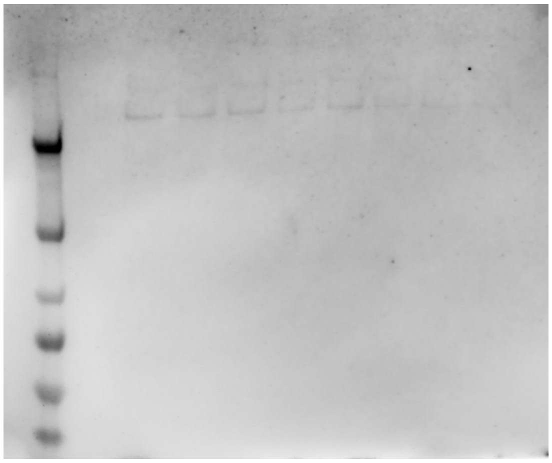

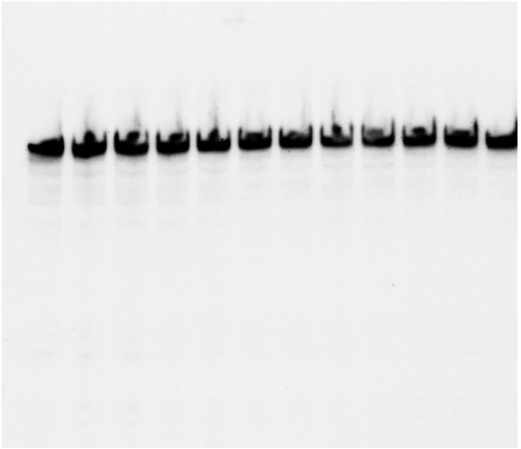

**Full Blots for Fig. S5 (All CRBN-binding Compounds)**

H3K18Ac

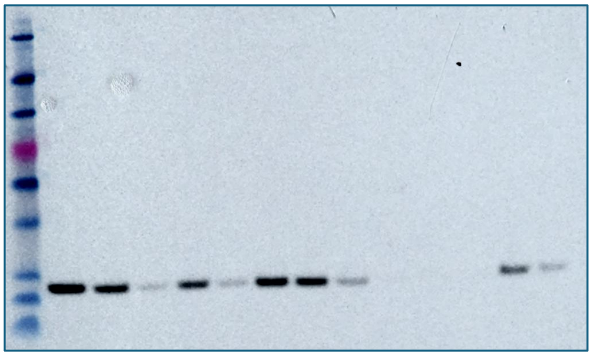

H3K23Ac

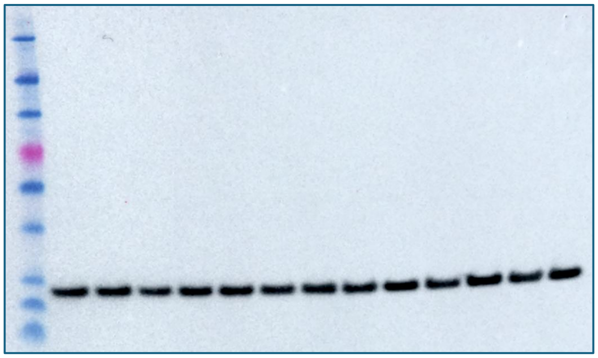

Ponceau

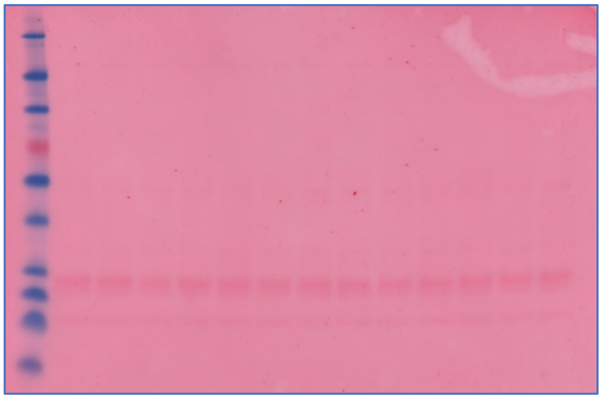

Ponceau

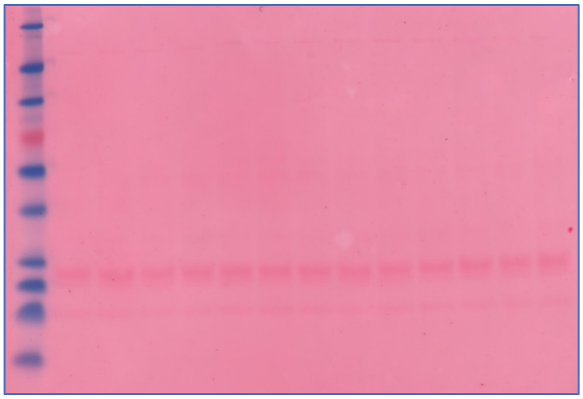

**Full Blots for Fig. S5 (All VHL-binding Compounds)**

H3K18Ac

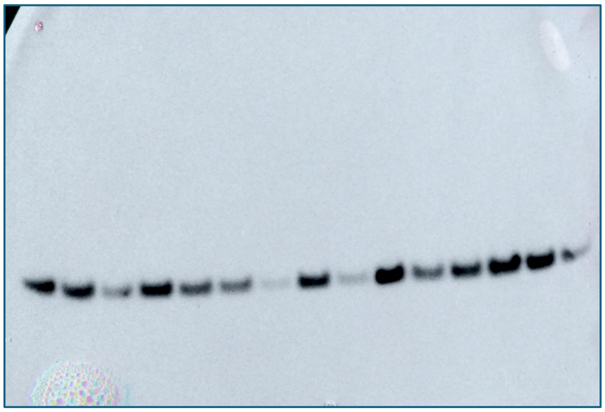

H3K23Ac

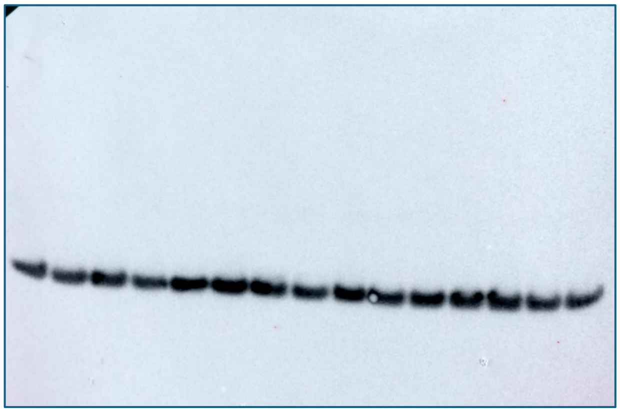

Ponceau

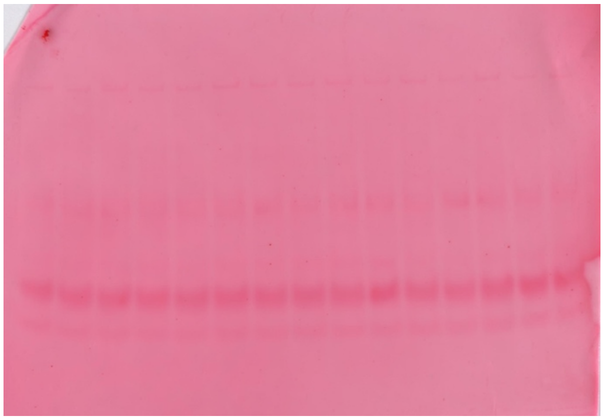

Ponceau

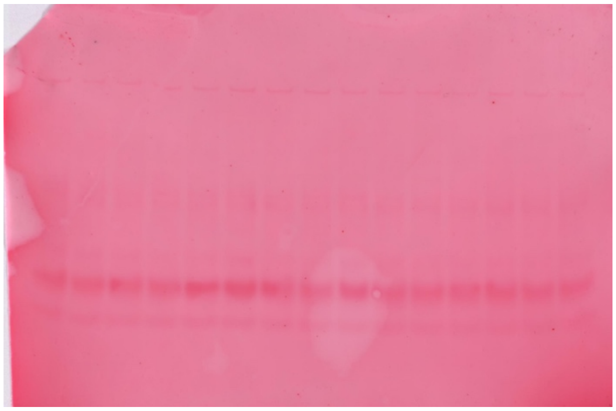
